# Novel Methods for Elucidating Modality Importance in Multimodal Electrophysiology Classifiers

**DOI:** 10.1101/2022.01.01.474276

**Authors:** Charles A. Ellis, Mohammad S.E. Sendi, Rongen Zhang, Darwin A. Carbajal, May D. Wang, Robyn L. Miller, Vince D. Calhoun

## Abstract

Multimodal classification is increasingly common in biomedical informatics studies. Many such studies use deep learning classifiers with raw data, which makes explainability difficult. As such, only a few studies have applied explainability methods, and new methods are needed. In this study, we propose sleep stage classification as a testbed for method development and train a convolutional neural network with electroencephalogram (EEG), electrooculogram, and electromyogram data. We then present a global approach that is uniquely adapted for electrophysiology analysis. We further present two local approaches that can identify subject-level differences in explanations that would be obscured by global methods and that can provide insight into the effects of clinical and demographic variables upon the patterns learned by the classifier. We find that EEG is globally the most important modality for all sleep stages, except non-rapid eye movement stage 1 and that local subject-level differences in importance arise. We further show that sex, followed by medication and age had significant effects upon the patterns learned by the classifier. Our novel methods enhance explainability for the growing field of multimodal classification, provide avenues for the advancement of personalized medicine, and yield novel insights into the effects of demographic and clinical variables upon classifiers.

## Introduction

In recent years, more biomedical informatics studies have begun to incorporate multimodal data into the training of machine learning and deep learning classifiers (1–3). The use of complementary modalities can enable the extraction of better features and improve classification performance (1)(4). While multimodal data can improve classifier performance, it can also make explaining models more challenging. This is especially true for state-of-the-art deep learning models that use automated feature extraction. As a result, most studies have not used explainability (4–8)(9), which is concerning because transparency is increasingly demanded for model development and physician decision making (10). As such, more multimodal explainability methods need to be developed (2,3,11–14). In this study, we propose the use of automated sleep stage classification as a testbed for the development of multimodal explainability methods. We further present 3 novel approaches for use with multimodal electrophysiology data that offer significant improvements over existing approaches. Specifically, we present a global ablation approach that is uniquely adapted for electrophysiology data. We further present two local methods that can be used to identify personalized electrophysiology biomarkers that would be obscured by global methods. Using the local methods, we then perform a novel analysis that illuminates the effects of demographic and clinical variables upon the patterns learned by the classifier.

### Automated Sleep Stage Classification as Testbed for Multimodal Explainability

Automated sleep stage classification offers a unique testbed for the development of novel multimodal explainability methods. Automated sleep stage classification has multiple noteworthy characteristics. (1) In practice, clinicians rely on multiple modalities instead of a single modality to manually score sleep stages (15). (2) The features differentiating sleep stages and the importance of modalities are well-characterized in a clinical setting (15). (3) Multiple large sleep stage datasets are publicly available (16–18). (4) A number of studies involving unimodal and multimodal sleep stage classification have been conducted (4)(19), which could enable data scientists to develop their explainability methods alongside established architectures. Because these characteristics can help us validate our explainability methods and because there is a clinical need for explainability in sleep stage classification, we chose sleep stage classification as a use-case in this study. In the following paragraphs, we first discuss the different stages of sleep. We then briefly review the domain of sleep stage classification and the explainability methods that have been used within the domain, both for multimodal and unimodal classification studies.

### Characteristic Features of Sleep Stages

Typical sleep stage classification approaches involve the classification of 5 stages: Awake, rapid eye movement (REM), non-REM1 (NREM1), NREM2, and NREM3. Clinicians use the American Academy of Sleep Medicine (AASM) manual (15) to identify the stages, and the stages are clearly described in other sleep staging studies (20)(21). According to the AASM manual (15), Awake periods are characterized by α band (8 – 13 Hz) electroencephalogram (EEG) activity in occipital brain regions when a participant’s eyes are closed. When a participant’s eyes are open, Awake periods are characterized by eye blinks, rapid eye movements, reading eye movements (i.e., slow movements followed by a burst of movement in the opposite direction), and strong levels of electromyogram (EMG) activity. NREM1 periods are characterized by slow eye movements, θ band (4 – 7 Hz) EEG activity, and vertex sharp waves (i.e., large V-shaped waves with a duration of less than 0.5 seconds). They have EMG activity at lower levels than Awake periods. NREM2 is principally characterized by EEG activity. It has K-complexes (i.e., sharp negative decreases in signal amplitude followed immediately by sharp increases in positive signal amplitude) and sleep spindles (i.e., trains of waves between 11 – 16 Hz). NREM2 typically has no electrooculogram (EOG) activity. It typically has less EMG than Awake periods and can have levels as low as REM. NREM3 mainly consists of d (0.5 – 2 Hz) EEG activity. NREM3 typically has little or no EOG activity. It has less EMG activity than NREM2 and can have EMG activity as low as REM. REM is characterized by rapid eye movements in the EOG, periods of little or no EMG activity followed by brief irregular bursts of activity, and sharp triangular EEG activity. Each stage is characterized by distinct EEG, EOG, and EMG activity.

### Unimodal Sleep Stage Classification and Explainability

Many sleep stage classification studies have used unimodal EEG. While some studies have used extracted features for sleep stage classification (19,20,22,23), recent studies have begun to use deep learning techniques involving automated feature extraction from raw data (21,22,24–27). Multiple studies have used explainability methods for unimodal classification. Some studies have used extracted spectral features. In a couple of studies, authors trained convolutional neural networks (CNNs) to classify EEG spectrograms and applied sensitivity or activation maximization (28) to identify the important features (29)(30). In other studies, authors trained interpretable machine learning models or deep learning models with layer-wise relevance propagation (LRP) (31) to classify power spectral density values and gain insight into the features learned by the classifiers (32)(33). A few studies involving deep learning models with raw data have also used explainability methods (25,34–36). These studies typically seek to identify the spectral features (34–39) or waveforms (36,37) learned by neural networks. However, multimodal classification poses unique challenges for explainability that do not exist for unimodal classification.

### Multimodal Explainability in Sleep Stage Classification and Other Domains

As such, most multimodal classification studies, regardless of whether they used extracted features (5,8) or deep learning-based automated feature extraction (4,7), have not used explainability methods. Among the few studies using explainability methods (40–42), some have used extracted features and forward feature selection (FFS) (42). Others have used raw data and ablation for insight into modality importance (41). Additionally, some have shown the importance of EEG spectra or performance increases after retraining a model with additional modalities (40). Some multimodal explainability methods have also been presented in other domains (2)(3)(43). Similar to (42), one paper used FFS to find key features from clinical scales and imaging features (3). One study used impurity and ablation (2). Another multimodal study showed the importance of parts of inputs for one modality (43) with Grad-CAM (44).

### Current Multimodal Explainability Methods

Multiple methods have been used for insight into modality importance in multimodal classifiers: FFS (3)(42), impurity (2), and ablation (2)(41). FFS is applicable to most classifiers. However, it requires repeated retraining, which is impractical for computationally intensive deep learning frameworks. Impurity is only applicable to tree-based classifiers. Lastly, ablation is, like FFS, also applicable to nearly any classifier and is easy to implement. In contrast, to FFS, ablation is not computationally intensive. As such, of existing approaches, it is most useful for finding modality importance in deep learning classifiers.

### Limitations of Ablation and Novel Alternative

Ablation does have a key weakness like all perturbation-based post-hoc explainability methods. Specifically, perturbation methods can create out-of-distribution samples that lead to a poor assessment of modality importance (45). Ablation also involves (1) the substitution of a feature or modality with values that are theoretically neutral (i.e., that do not give evidence for a particular class) and (2) an examination of how that ablation affects the classifier. As such, when translating ablation to a new domain, it is important to consider how to set a modality or feature to a neutral state while minimizing the likelihood of creating out-of-distribution samples or features. Existing studies using ablation for insight into multimodal classifiers have replaced each modality with zeros (2)(41). However, zeroing out modalities creates samples that are highly irregular within the electrophysiology domain. The only instance in which an electrode might return a value of zero is when the electrode is completely disconnected from the device. In contrast, electrodes commonly return line-related noise or, in instances when an electrode is not working properly, only return line-related noise. Line-related noise is found in electrophysiology data at 50Hz or 60Hz due to the presence of lights, power lines and other electronics near recording devices. Because it is so often found in electrophysiology data, a classifier should learn to ignore it, and it should be neutral to the classifier. As such, line-related noise could offer a more reliable, electrophysiology-specific alternative to the generic zero-out ablation methods that have previously been applied.

While line noise-based ablation would be less likely to produce out-of-distribution samples or features than a typical zero-out ablation approach, it would still be at risk of doing so. Gradient-based feature attribution (GBFA) methods (46) offer an alternative to ablation that does not risk producing out-of-distribution samples. GBFA methods include approaches like LRP (31), Grad-CAM (44), and sensitivity (28).

### Limitations of Global Explanations and Proposal of Novel Local Explainability Approach

Global explainability methods identify the overall importance of each feature or modality to the classifier. In contrast, local methods provide higher resolution insight and indicate the importance of each feature or modality to the classification of individual samples (45). Global methods have inherent limitations relative to local methods, and existing multimodal explainability approaches have mainly been global. Importantly, as shown in Figure 1, global explanations obscure feature importance for individual samples and can obscure the presence of subgroups. Local explanations for many samples can be combined for higher level or global importance estimates (33) (34). Because of this, they can also be analyzed on a subject-specific level that paves the way for the identification of personalized biomarkers. Furthermore, local explanations can be used to examine the degree to which demographic and clinical variables affect the patterns learned by a classifier for specific classes and features (13), which is a capacity that has not previously been exploited in multimodal classification. Local methods have been applied in a couple multimodal classification studies. In one study, authors ablated time points of an input sample and examined the effect on the classification of the sample (41). In another study, authors used Grad-CAM to examine segments of a single modality (43). Neither study identified the importance of each modality.

**Figure 1.**
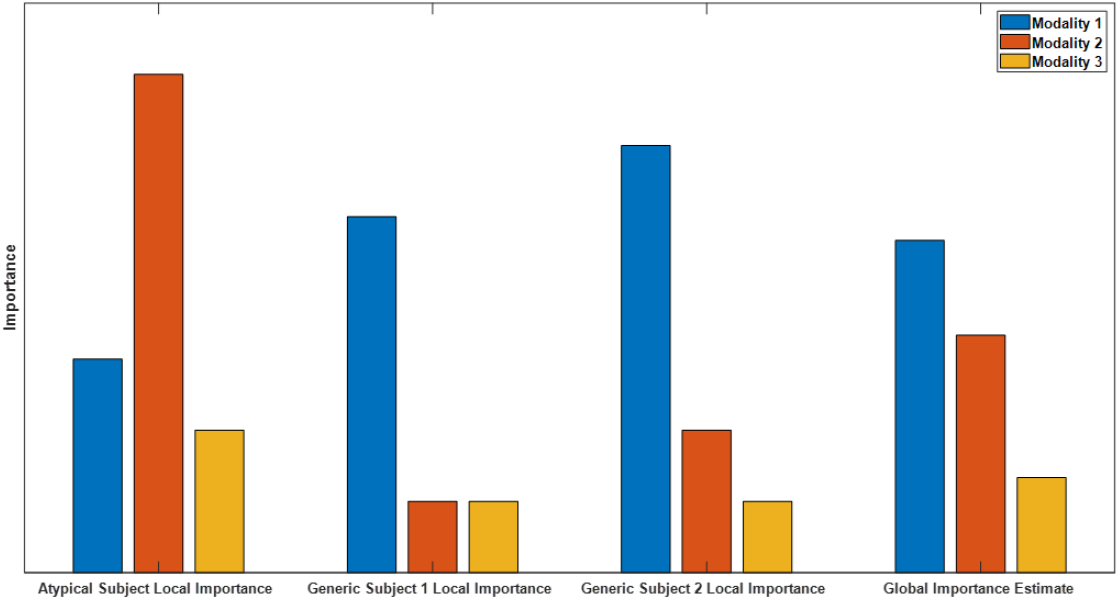
Example of Global versus Local Importance with Dummy Data. From left to right, are 4 importance metrics: the importance for an atypical sample, the importance for two generic samples, and the global importance estimate that is formed by averaging the importance values for the individual samples. Note that modality two is very important for the atypical subject but not for the generic subject, so the presence of the atypical sample is hidden in the global importance. When thousands of samples from dozens of subjects are being analyzed, the presence of subgroups is easily obscured.

In the present study, we train a CNN for automated sleep stage classification using a popular publicly available dataset. We introduce a novel global ablation approach that is uniquely adapted for the electrophysiology domain (12). We then present a novel local ablation approach (13) and show how GBFA methods can be used for local insight into multimodal classifiers (11). With our local methods, we identify subject-level differences in modality importance that support the viability of the methods for the identification of personalized biomarkers. We then use the local explanations to perform a novel analysis that provides insight into the patterns learned by the classifier related to the age, sex, and state of medication of subjects in our sleep dataset (13,14).

## Results

Here, we describe our model performance, explainability, and statistical analysis results.

### Model Performance Results

Table 1 shows the mean and standard deviation of the precision, recall, and F1 score for each class. The model had highest F1 scores for NREM2 and Awake. Possibly because of its smaller sample size, NREM1 had the lowest classification performance across all metrics. While performance for NREM3 and REM was not as high as for NREM2 and Awake for most metrics, the classifier still performed well for both classes.

**Table 1.** Classification Performance Results

|  | Awake | NREM1 | NREM2 | NREM3 | REM |
| --- | --- | --- | --- | --- | --- |
| **F1** | 71.25±05.15 | 39.86±07.19 | 73.28±04.76 | 64.15±15.25 | 65.92±06.28 |
| **Precision** | 72.25±07.12 | 36.20±03.98 | 79.35±03.92 | 56.78±18.35 | 69.04±07.14 |
| **Recall** | 70.90±07.02 | 46.28±13.52 | 68.71±08.51 | 78.22±10.24 | 63.26±06.69 |

Table 2 shows a confusion matrix of the classifier performance on test samples across all folds. A relatively large number of Awake samples were assigned to the NREM1 class, and NREM1 samples were assigned to all classes but NREM3. Many NREM2 samples were assigned to NREM3, and many NREM3 samples were assigned to NREM2. Many REM samples were assigned to NREM1 and NREM2.

**Table 2.** Confusion Matrix for Test Samples Across All Folds

|  |  | Predicted |  |  |  |  |  |
| --- | --- | --- | --- | --- | --- | --- | --- |
|  |  | Awake | NREM1 | NREM2 | NREM3 | REM | Totals |
| Actual | Awake | 4574 | 1083 | 293 | 151 | 240 | 6341 |
|  | NREM1 | 806 | 2424 | 879 | 212 | 890 | 5211 |
|  | NREM2 | 602 | 1981 | 17965 | 3928 | 1902 | 26378 |
|  | NREM3 | 133 | 108 | 1315 | 6125 | 74 | 7755 |
|  | REM | 169 | 1157 | 2197 | 472 | 6929 | 10924 |
|  | Totals | 6284 | 6753 | 22649 | 10888 | 10035 | 56609 |

### Global Ablation Results

Figure 2 and Supplementary Figure 1 show the results comparing our noise-related global ablation analysis with the typical global ablation approach that zeroes out a modality for correct classification groups and all classification groups, respectively. We computed the percent change in the number of samples assigned to a classification group within the confusion matrix after perturbation. Figure 2 shows bar plots of the median importance for each group, and Supplementary Figure 1 shows box plots for each group.

**Figure 2.**
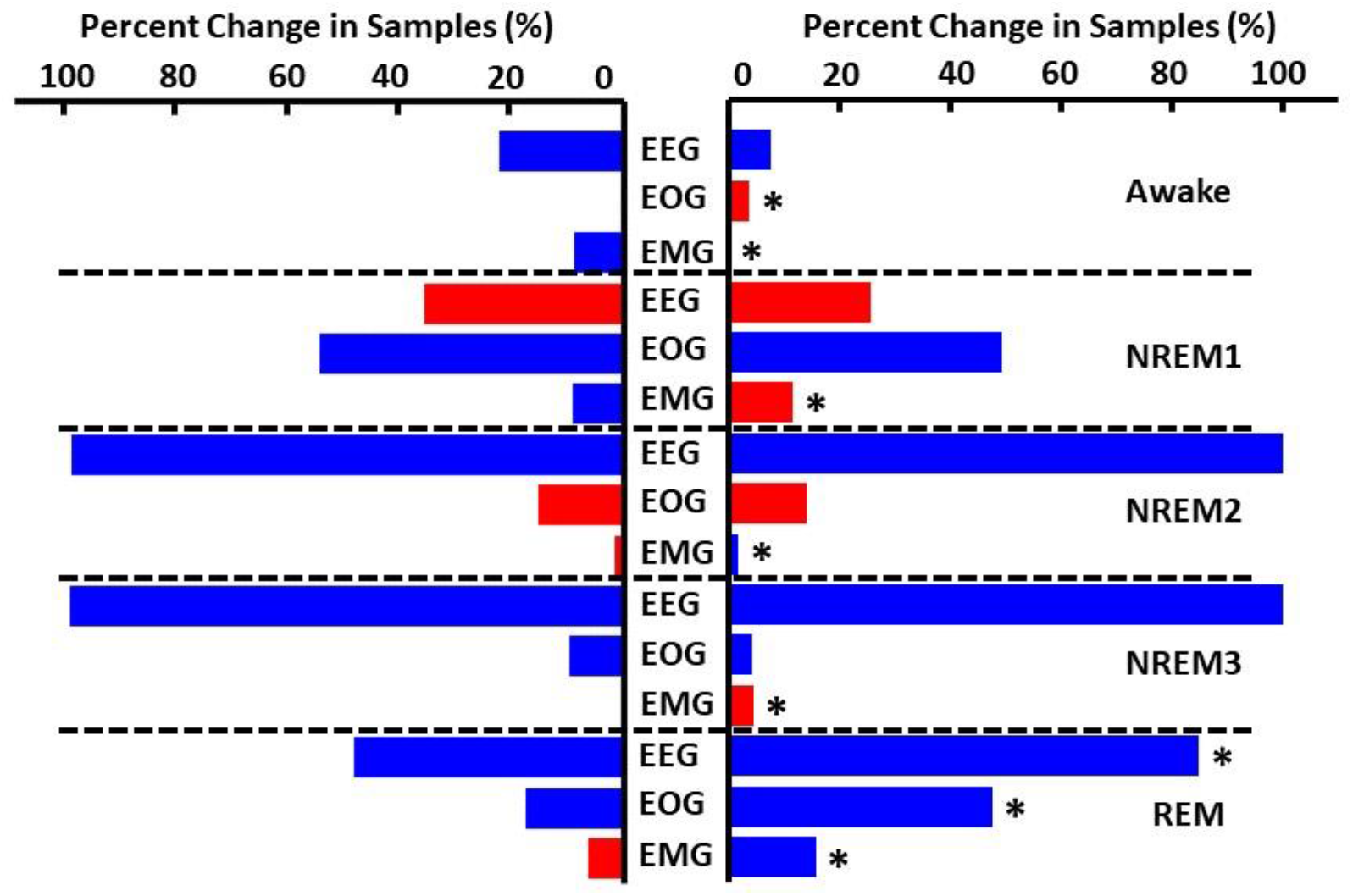
Global Ablation Results for Correctly Classified Samples. The leftmost and rightmost plots show the results for the zero-out ablation and our noise-related ablation, respectively. Importance for each sleep stage and modality is aligned vertically. Horizontal dashed lines separate importance for each sleep stage. Blue and red bars indicate negative and positive percent change in samples, respectively. Asterisks indicate significant differences (p < 0.05) between results for the two methods. Across both methods, EEG was most important for all classes except NREM1.

In this paragraph we discuss correctly classified samples. EEG was the most important modality across both methods for all classes except NREM1. For Awake samples, the model was most affected by EEG ablation, though the effects of ablating EEG varied widely across folds. Interestingly, the two methods had significantly different effects on the change in the number of Awake samples following EOG ablation. For EOG, our approach caused the number of Awake samples to increase, while the typical approach caused the number of Awake samples to decrease. The two methods also significantly differed for EMG. Ablating EMG with the typical approach decreased the number of Awake samples, while ablating EMG with our approach did not really have an effect. For NREM1, EEG and EOG ablation had large effects for both methods. EEG ablation increased the number of NREM1 samples, and EOG ablation decreased the number of NREM1 samples for both standard and noise-related methods. NREM1 EMG results significantly differed for the two methods. Ablation of EMG with our approach increased the number of NREM1 samples, and ablation with the typical approach slightly decreased the number of NREM1 samples. For NREM2, ablation of EEG caused nearly all NREM2 samples to be misclassified across nearly all folds. Ablation of EOG increased the number of NREM2 samples, and ablation of EMG had little effect. For NREM3, ablation of EEG caused nearly all NREM3 samples to be misclassified. EOG ablation caused an increase in misclassified NREM3 samples. EMG ablation had little effect, though our ablation approach showed a slight increase in NREM3 samples in some instances. For REM, EEG and EOG ablation caused large reductions in REM samples. In both instances, ablation with our approach produced a significantly larger reduction than ablation with the typical approach. Also, EMG ablation with the standard approach had negligible effects but a significant decrease in REM samples with our approach.

Because we measured the percent change in samples assigned to each classification group, the effects of perturbing modalities to different classification groups can be compared. Ablating NREM2 and NREM3 EEG prompted a much larger decrease in correctly classified samples than ablating EEG for other classes, though the effect on REM was only slightly less. In contrast, ablating EEG significantly increased the number of correctly classified NREM1 samples. EOG ablation had the strongest impact upon NREM1 followed by REM, where the number of correctly classified samples decreased. EMG seemed to have little effect upon the classifier. However, EMG did impact Awake and REM classification to a small degree.

For samples misclassified as Awake, EEG ablation greatly increased the number of samples from other classes that were assigned to Awake. EMG and EOG ablation only affected the number of NREM1 samples assigned to Awake (i.e., NREM1/Awake), though they affected the number of REM/Awake to a smaller degree. For samples misclassified as NREM1, EEG ablation significantly increased the number of misclassified samples. For samples misclassified as NREM2, EEG ablation prevented samples from most classes being classified as NREM2, but EOG ablation increased the number of misclassified samples. EEG and EOG ablation also reduced the number of samples misclassified as NREM3. Ablation had negligible effects upon the number of NREM2/REM and NREM3/REM. Interestingly, the two ablation methods seemed to have approximately opposite effects for the number of Awake/REM and NREM1/REM.

### Local Ablation Results

We performed several analyses with our local ablation approach. (1) We combined the results across samples to form a global estimate of modality importance. (2) We showed how the local explanations evolved over a 2-hour window of data. (3) We examined how the ablation results might give insight into the patterns learned by the classifier related to demographic and clinical variables.

The bar plots show the results for the correct classification group. EEG, EOG, and EMG importance values are shown in red, green, and blue, respectively. EEG was most important for all classes except NREM 1.

#### Comparison of Global Ablation with Local Ablation-based Global Estimation of Modality Importance

Figure 3 and Supplementary Figure 2 show how we applied our local ablation approach across all samples in each fold to estimate global modality importance for correct classification groups and all classification groups, respectively. This enabled us to validate our local ablation approach by comparing it to our global ablation approach. To find the effects across samples, we calculated the absolute percent change for each sample and then calculated the mean value across samples. Importantly, like in global ablation, our local ablation approach found that EEG was the most important modality for all classes except NREM1 in which EOG was most important. Overall, our local and global ablation approaches seemed to assign similar levels of relative importance. However, this was not true for many incorrect classification groups. For example, the relative importance of each modality differed across methods for NREM3/NREM2 and REM/NREM2.

**Figure 3.**
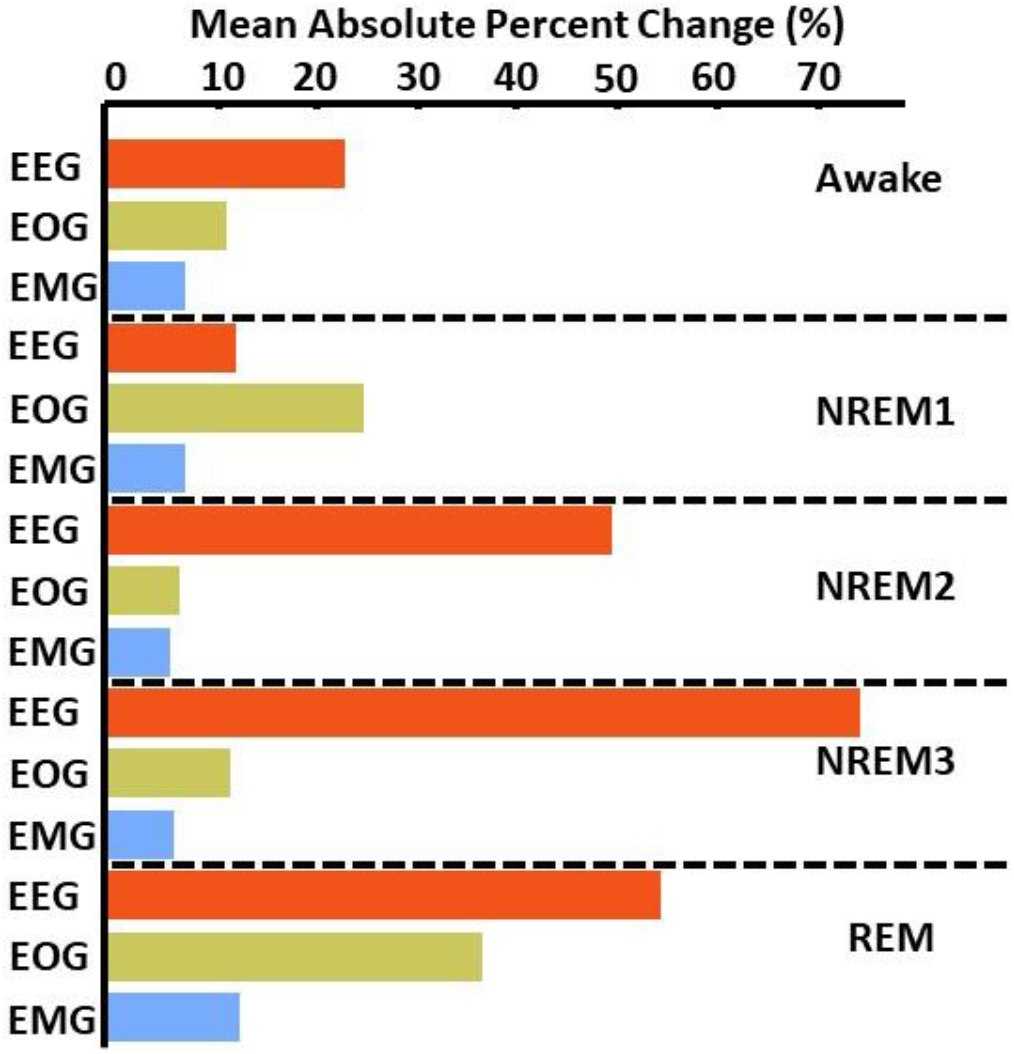
Local Ablation Results Showing Estimation of Global Importance for Correct Classification Groups. Within each fold, we calculated the mean absolute percent change in activation for the perturbation of samples in each classification group. We then calculated the median value across folds.

#### Subject-Level Local Ablation Results Over Time

Figure 4 shows the local ablation results over the first two hours of a recording from Subject 12. Note that the 2-hour period is roughly equivalent to a full sleep cycle, as the period starts while the subject is awake and cycles through all sleep stages. Importantly, EOG was often more important during Awake periods than EOG. Interestingly, an EMG spike occurred in the first five minutes of the recording that increased the likelihood of samples being classified as Awake when ablated. In contrast, when EEG and EOG were ablated, the likelihood of the samples being classified as Awake decreased. As Subject 12 proceeded into NREM1 at around 20 minutes, EOG ablation significantly decreased the likelihood of the samples being classified correctly. EEG and EMG ablation decreased the likelihood of the samples being correctly classified at the beginning and ending of the period and increased the likelihood of the samples being correctly classified during the middle of the period. As Subject 12 went into deeper sleep stages, EEG ablation tended to decrease the probability that a sample would be classified as NREM. Additionally, EOG importance varied greatly throughout the remaining time course.

**Figure 4.**
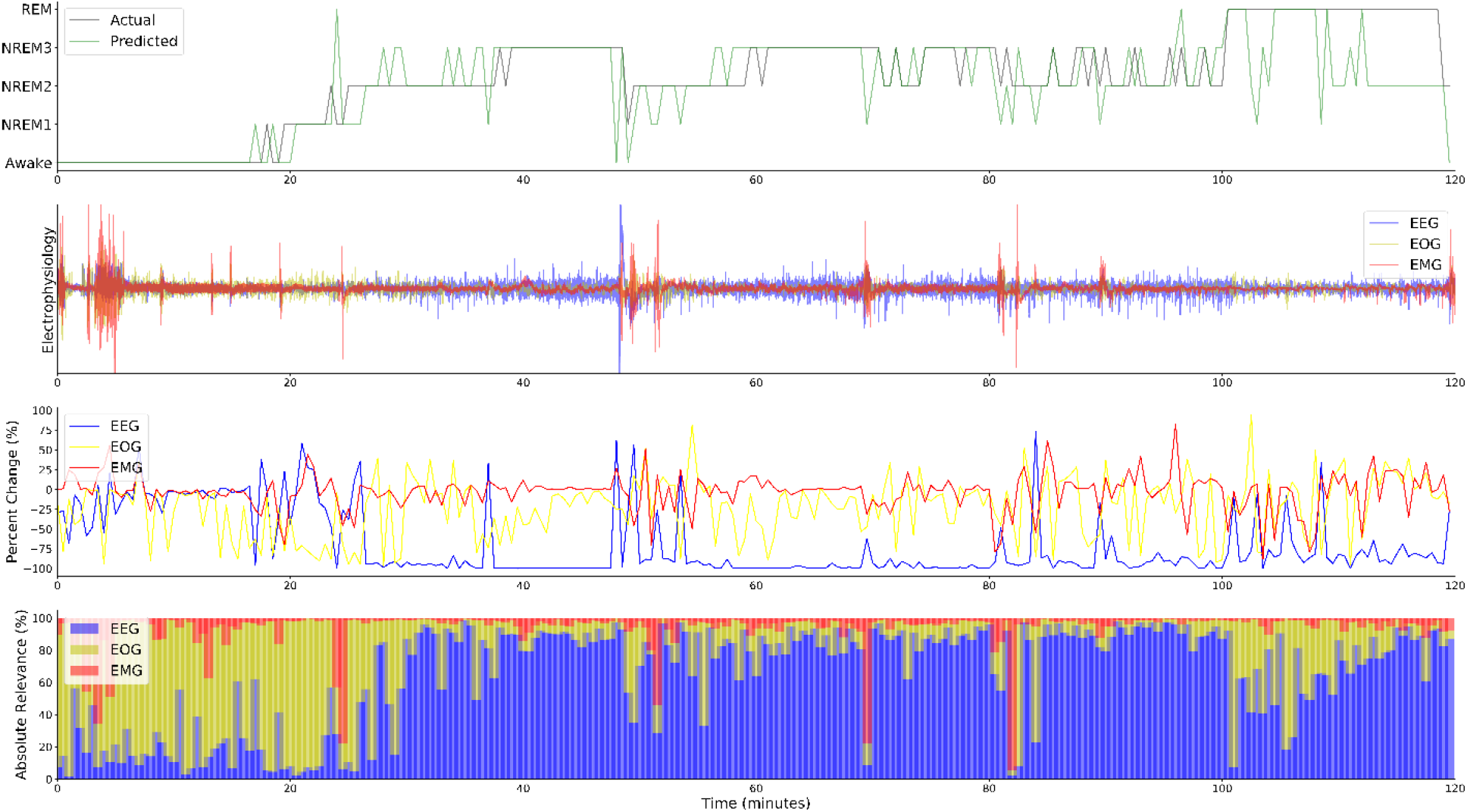
Local Explanations over a 2-Hour Sleep Cycle from Subject 12. The top panel shows the actual and predicted classes. The second panel shows electrophysiology activity. The third panel shows the local ablation results, and the fourth panel shows the percent of LRP relevance (ε = 100) for each modality. In contrast to the global results, EOG is more important for Awake than EEG from 0 to 20 minutes.

#### Statistical Analysis of Local Ablation Results with Demographic and Clinical Variables

Figure 5 and Supplementary Figure 3 show the results for the statistical analysis examining the effects of medication, sex, and age upon the local ablation explanations for correct classification groups and all groups, respectively. The importance assigned to many of the modalities and classification groups had significant relationships with the demographic and clinical variables. Importantly, subject sex had significant relationships with 13 of the 15 correct classification group modality pairs, while medication and age only had relationships with 10 and 9 of the pairs, respectively.

**Figure 5.**
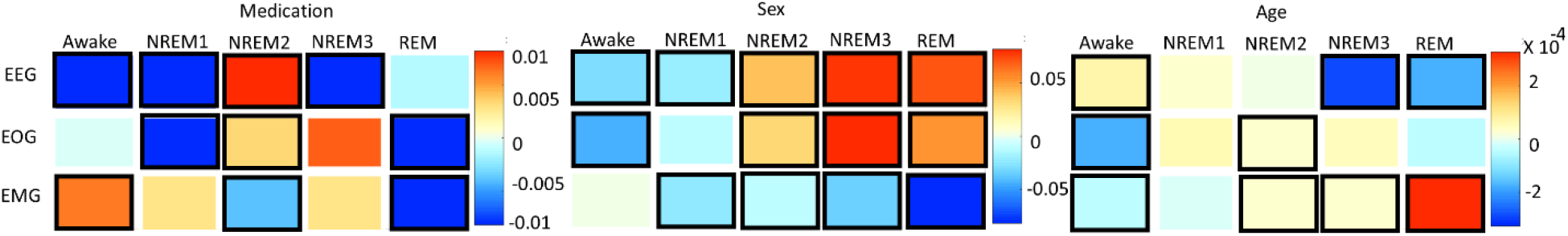
Effects of Clinical and Demographic Variables upon Local Ablation Importance for Correct Classification Groups. Panels a, b, and c show results for medication, sex, and age, respectively. The x -axes indicate the predicted class, and the y-axis indicates the modality. The heatmaps show the regression coefficient values. Squares surrounded by a bolded box show significant effects (p < 0.05). A positive medication coefficient indicates that temazepam samples had more importance than placebo samples. For subject sex, a positive coefficient indicates that female samples had more importance than male samples, and a positive age coefficient indicates that importance increased with age. Note that sex had more significant relationships than the other variables.

More EOG groups had significant relationships with medication than EEG or EMG groups. For EEG, the explanations for all correctly classified groups, except REM, had significant relationships with medication. This is interesting when one notes that for REM/REM samples EOG and EMG importance had significant relationships with medication. The strongest EEG relationships were for NREM1/NREM3 and NREM1/REM. Interestingly, the importance of most samples assigned to NREM3 had significant relationships with medication. For EOG, most NREM1 and NREM2 groups had significant relationships with medication. The only instance in which importance increased from placebo to temazepam was for the NREM2/REM. For EMG, NREM2 importance had relationships with medication for all groups except Awake. Additionally, NREM3/NREM2 and REM/NREM2 also had significant relationships with medication.

Subject sex was related to many classification groups across modalities. For EEG, all correct classification groups had significant relationships with subject sex. Additionally, samples predicted to be NREM3 had very strong positive relationships with sex. Samples belonging to female subjects had higher importance for samples misclassified as NREM3. Samples belonging to male subjects seemed to have higher EEG importance for REM/Awake. Interestingly, EEG appeared to be more salient in samples belonging to female subjects than male subjects while the opposite was true for EMG. EOG groups had both positive and negative coefficient values. Samples belonging to female subjects that were misclassified as Awake had less EOG importance than those belonging to male subjects. In contrast, samples belonging to female subjects that were misclassified as NREM2 had more importance than those belonging to male subjects. The EOG coefficients with the largest magnitude were all associated with NREM3. The largest magnitude EMG coefficients were for samples classified as REM, and particularly Awake/REM. Male subjects had more EMG importance for Awake and NREM1 samples than female subjects.

For subject age, the classifier put more importance upon EEG in Awake as age increased. Also, except for REM/Awake, the classifier placed less importance on EEG in REM as age increased. REM/Awake and Awake/NREM3 had more EEG importance with increased age, and NREM3/Awake and REM/NREM3 had less EEG importance with increased age. EOG importance for NREM2 samples was related to modest increases in age. Interestingly, correctly classified Awake and NREM1/REM had significant decreases in EOG importance with increased age. REM/NREM3 had significantly increased EOG importance in samples with increased age. Most correct classification groups had a relationship between the EMG importance and subject age. Except for correctly classified Awake samples, EMG importance generally increased with age. Misclassified Awake and REM samples also had strong relationships between EMG importance and age. NREM3/Awake and REM/Awake had more EMG importance with age, though NREM2/Awake had less EMG importance with age. NREM1/REM had less EMG importance with increased age.

### LRP Results

We performed several analyses upon the LRP relevance. (1) For a global estimate of modality importance, we computed the percentage of absolute relevance assigned to each modality across folds for each classification group. (2) We showed the local results for the same 2-hour window as the ablation analysis. (3) We examined relationships between absolute LRP relevance and clinical and demographic variables.

#### Global Estimation of Modality Importance with LRP

Figure 6 and Supplementary Figure 4 show the LRP results for correct classification groups and all groups, respectively. EEG was more important or of comparable importance to EOG across most classes. For correctly classified Awake, NREM1, and REM, EEG and EOG were of comparable importance for the ε-rule (ε = 100) and the αβ-rule. However, EEG was more important than EOG for the ε-rule (ε = 0.01). For NREM2/NREM2 and NREM3/NREM3, EEG was more important than EOG and EMG. EMG importance was greatest in Awake, followed by NREM2 and NREM1. Relative to correctly classified groups, many incorrectly classified groups had greater variance in relevance across folds. Nevertheless, samples incorrectly assigned to a class typically had similar relevance distributions to samples correctly assigned to the class. NREM2, NREM3, and REM assigned to Awake were exceptions to this pattern. They demonstrated high levels of variance across folds. NREM2 and NREM3 incorrectly classified as REM and NREM2 also demonstrated increased variance across folds relative to REM.

**Figure 6.**
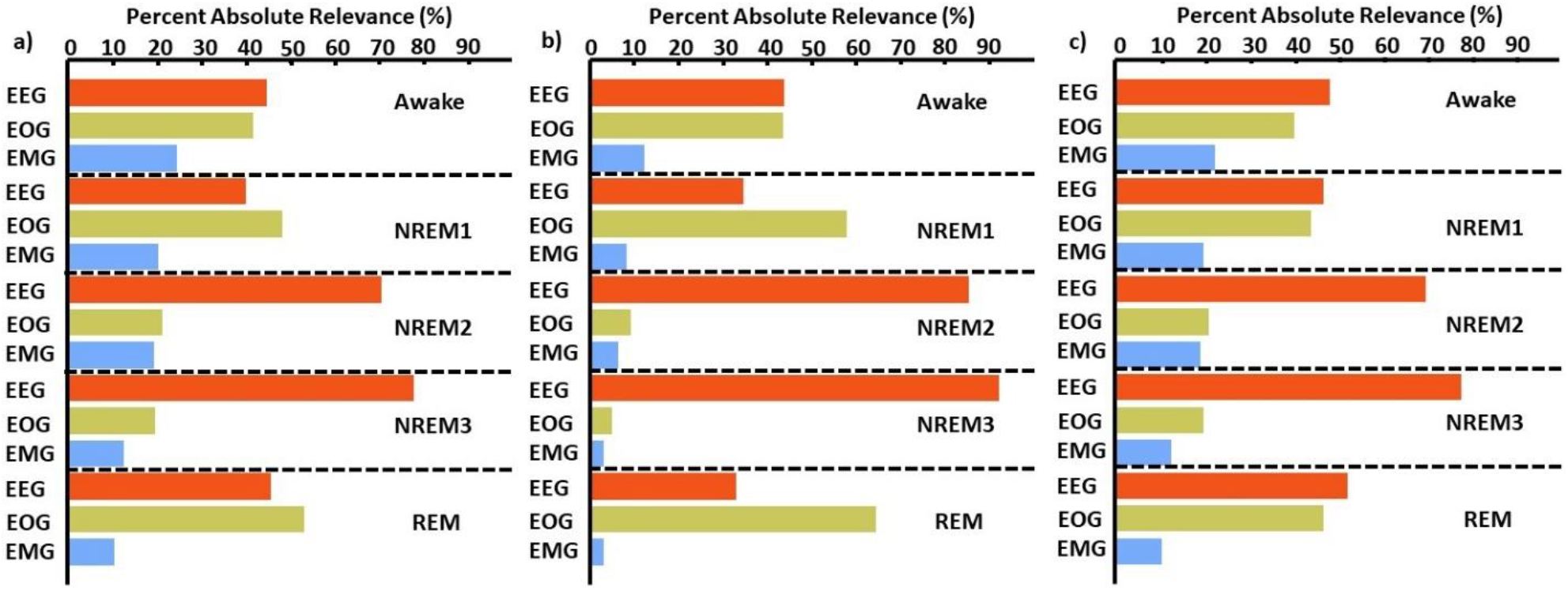
LRP Global Estimate of Modality Importance for Correct Classification Groups. We calculated the percent of absolute relevance for each modality across samples within each classification group. We then calculated the median value across folds. Panels a, b, and c show results for the αβ-rule, ε-rule (ε =100), and ε-rule (ε=0.01), respectively. Red, green, and blue bars are for EEG, EOG, and EMG, respectively. EEG was, with a few exceptions, most important.

#### LRP Results Over Time

The bottom panel of Figure 3 shows the local LRP results for the ε-rule (ε = 100) over the same 2-hour period for which the local ablation analysis was conducted. During the first 17 minutes, the model correctly classified Awake by heavily relying upon EOG. EOG also demonstrated high levels of importance for REM between 100 to 120 minutes. For correctly classified NREM1 through NREM3, the model heavily relied upon EEG. However, when NREM1 through NREM3 were misclassified, the model often placed greater importance upon EMG and EOG (i.e., 70 to 80 minutes). EMG overall had little relevance. However, spikes in EMG relevance corresponded with spikes in EMG activity throughout the recording.

#### Statistical Analysis of Local LRP Results with Demographic and Clinical Variables

Figure 7 and Supplementary Figure 5 show the results of our analyses examining the relationship between LRP relevance and medication, sex, and age for correct classification groups and all groups, respectively. Significant relationships existed for many of the modalities and classification groups. Importantly, subject sex had significant relationships with 14 of the 15 correct classification group modality pairs, while medication and age only had relationships with 11 of the pairs.

**Figure 7.**
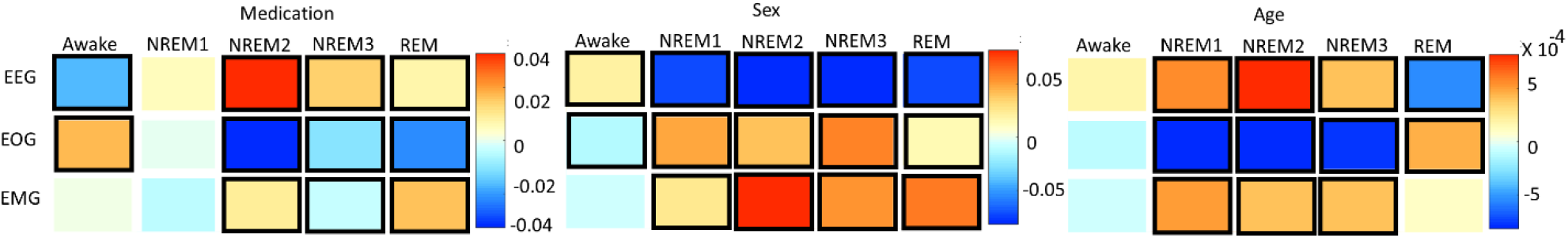
Effects of Clinical and Demographic Variables upon Local LRP Relevance for Correct Classification Groups. Panels a, b, and c show results for medication, sex, and age, respectively. The x-axis of each panel indicates the predicted class, and the y-axis indicates the modality. The heatmaps show the size of the regression coefficient. Squares surrounded by a bolded box show significant effects (p < 0.05). For medication, a positive coefficient indicates that temazepam samples had more importance than placebo samples. For subject sex, a positive coefficient indicates that female samples had more importance than male samples, and for age, a positive coefficient indicates that importance increased with age. Note that sex had more significant relationships than the other variables.

For medication, EEG had more relevance across many groups for temazepam relative to placebo samples, while EOG had a decrease in relevance and EMG had mixed results. For EEG, the only group with decreased relevance for temazepam relative to placebo was Awake/Awake, which corresponded with increased relevance for temazepam relative to placebo in EOG. For EEG, all correct classification groups except NREM1 had significant relationships with medication. For incorrect classification groups, NREM1/NREM3 had the highest magnitude coefficient. This corresponded with a large decrease in EOG relevance. For EOG, placebo samples assigned to NREM2 and NREM3 had less relevance than temazepam samples. NREM2/Awake also had a strong relationship with medication. For EMG, fewer classification groups had significant relationships with medication than for EEG and EOG. However, correctly classified NREM2, NREM3, and REM had significant relationships with medication. Awake/NREM3 had a large coefficient, with temazepam samples having more EMG relevance than placebo samples.

For subject sex, EEG had more importance in males than females across most classification groups. In contrast, EOG and EMG had more relevance for females than males across most groups. For EEG and EOG, all correct classification groups were related to subject sex. For EMG, all correct classification groups except for Awake had significant relationships with sex. For EEG, the highest magnitude coefficients were REM/NREM2, REM/NREM3, NREM1/NREM3, and NREM3/REM. Male subjects had more EEG relevance than female subjects for REM/NREM2. Female subjects had more EEG relevance than male subjects for NREM1/NREM3. Interestingly, NREM3/REM and REM/NREM3 had coefficients of a nearly opposite values. For EOG, Awake/NREM3, NREM3/NREM1, and REM/NREM3 had the largest magnitude relationships with subject sex. Male subjects had much higher EOG relevance for Awake/NREM3 and for REM/NREM3 relative to female subjects, and female subjects had much higher EOG relevance for NREM3/NREM1. Samples belonging to female subjects tended to have more EMG relevance for correct classification groups than male subjects. The largest magnitude EMG coefficients were NREM1/NREM3, NREM3/NREM1, REM/NREM2, and REM/NREM3. For male subjects, NREM1/NREM3 and NREM3/NREM1 tended to have more relevance than for female subjects. For female subjects, REM/NREM2 and REM/NREM3 had more and less EMG relevance relative to male subjects, respectively.

For age, EEG and EMG relevance tended to increase with age, while EOG relevance decreased with age. For EEG, most classification groups, except for many Awake groups, had significant relationships with age. The largest magnitude EEG coefficients were for NREM3/Awake and REM/NREM3 where relevance decreased with age. For REM/NREM3, decreased EEG relevance with age corresponded to increased EMG relevance. For EOG, only REM/REM increased in relevance with age. This increase corresponded with decreased EEG relevance in REM/REM. For EMG, most groups with significant relationships with age had increased relevance with age. However, all classes misclassified as Awake that had significant relationships with age had a decrease in relevance with age. This corresponded to increased EEG relevance with age.

### Comparison of Explainability Results from Different Methods

To better understand our explainability methods, we compared the similarities and differences of their results. We compared the global importance estimates for global ablation, local ablation, and LRP. We also compared the local results over time and statistical analyses for local ablation and LRP.

#### Global Estimation

We compared the relative magnitude of the estimates across methods. Across methods, EEG was generally most important. For Awake/Awake, all three methods found that EEG was most important. However, LRP magnified the importance of EOG and EMG relative to EEG more than local ablation. For NREM1/NREM1, two LRP rules, local ablation, and global ablation found that EOG was most important, followed by EEG. Results for NREM2/NREM2 and NREM3/NREM3 were similar across explainability methods. EEG was most important, followed by EOG and EMG. For REM/REM, only the LRP ε-rule (ε = 0.1) agreed with local and global ablation regarding the relative modality importance. They identified the order of descending modality importance as EEG, EOG, and EMG. Many incorrect classification groups had similar distributions of relative importance across methods. However, some groups had different importance distributions. NREM1/Awake generally had greater EOG than EEG relevance for LRP but not for ablation. NREM2/NREM1 had less EOG than EEG relevance for LRP but not for ablation. Awake/REM and NREM1/REM had more EEG than EOG importance for local ablation but not for LRP. Global ablation found that EEG and EOG importance for Awake/REM varied according to the global ablation approach. Additionally, global ablation found that EEG had greater importance than EOG for NREM1/REM.

#### Subject-level Local Results over Time

In general, both local ablation and LRP showed similar trends in modality importance over time. They both showed lower levels of EEG and higher levels of EOG importance during Awake and NREM1 periods and showed a transition to higher EEG and lower EOG importance for NREM2, NREM3, and REM. However, in multiple instances, LRP seemed to more closely correspond with changes in electrophysiology activity. For example, between 60 and 80 minutes into the recording, EMG activity spiked, and a misclassification resulted. In this instance, LRP more clearly indicated that the change affected the classification than local ablation did. Additionally, for NREM periods between 30 to 100 minutes, EEG relevance often seemed to have greater variation relative to the relevance of other modalities than the local ablation results did.

#### Statistical Analysis of Effects of Clinical and Demographic Variables upon Local Explanations

Many effects were consistent between the two explainability methods. Importantly, across both methods, subject sex had relationships with more correct classification group modality pairs than either medication or age. The importance of EEG for Awake/Awake was less in temazepam than placebo samples and was more in temazepam than placebo samples for NREM2/NREM2. Samples assigned to NREM3 also generally had more EEG importance for temazepam than placebo samples. For EOG, most groups with significant relationships with medication had more importance across both methods in placebo than in temazepam samples. In REM/REM, EOG importance was higher in placebo than temazepam samples. NREM2/NREM1 and NREM3/NREM2 had more EMG importance in temazepam than placebo samples for both methods. REM/NREM2 had less EMG importance in temazepam than placebo samples for both methods.

For the effects of subject sex on EEG importance, the two methods provided similar results in many instances: (1) NREM1/NREM1, NREM1/NREM2, and NREM1/NREM3, (2) NREM2/NREM1, (3) REM/NREM3, and (4) Awake/REM. However, there were opposite effects in other cases: (1) for NREM2/NREM2, NREM2/NREM3, and NREM2/REM, (2) for NREM3/NREM3 and NREM3/NREM2, and (3) REM/REM and REM/NREM2. For EOG, correctly classified Awake, NREM2, NREM3, and REM had similar changes in importance from male to female samples. Additionally, the changes in importance for most classification groups were similar across methods. For EMG and sex, this was not the case, though in a few instances the differences in importance between male and female were similar (e.g., Awake/REM, NREM1/NREM2, NREM1/NREM3, and REM/NREM3).

Across explainability methods, subject age significantly affected the importance assigned to modalities. For EEG, there were similar effects of age: (1) for Awake/NREM3, (2) for NREM1/Awake and Awake/NREM2, (3) for NREM2/NREM1 and NREM2/REM, (4) for NREM3/Awake, and (5) for REM/REM, REM/Awake, and REM/NREM3. Many groups with different results across explainability methods had low levels of significance (i.e., p < 0.05). For EOG, there were differences in which classification groups were significant. However, NREM1/REM did have similar results across explainability methods. For EMG, there were many similarities in the effect of age upon the explanations of the two methods: (1) NREM1/NREM2, (2) NREM/NREM2 and NREM2/Awake, (3) NREM3/NREM3, and (4) REM/NREM3.

## Discussion

In this section, we discuss the implications of our methods to other multimodal classification problems. We then discuss the results that were uniform across explainability approaches within the context of established sleep domain knowledge, and we discuss the limitations and next steps for this work.

### Implications of Novel Explainability Methods Beyond Sleep Stage Classification

In this study, we present a series of novel multimodal explainability methods. Our global ablation method offers a methodological improvement over existing methods and is uniquely adapted to multimodal electrophysiology data. Existing multimodal electrophysiology studies have ablated each modality by replacing them with zeros (2)(41). Because having a channel of zero values is atypical for electrophysiology data in clinical settings, replacing modalities with zeros could increase the risk of creating out-of-distribution samples or features, which could cause the explanations to be inaccurate. Our approach adapts ablation for the domain of electrophysiology by replacing modalities with line-related noise that is commonly found in data and that the classifier should learn to ignore. Additionally, it highlights the usefulness of carefully considering the domain to which ablation is being adapted and using domain-specific perturbations. Our local ablation approach is, to the best of our knowledge, the first local multimodal explainability method that provides insight into the importance of each modality. By examining the change in output activation following ablation, it also shows how ablation or perturbation could be used to obtain local explanations across a variety of explainability problems beyond multimodal explainability. We also show, for the first time, how gradient-based methods can be used to find modality importance both locally and globally. Because they do not perturb data, GBFA methods could offer a more reliable approach than ablation. Our local methods offer a pathway towards identifying subject-specific electrophysiology biomarkers that could be used in personalized medicine. Additionally, our analysis of the relationship between the local explanations and demographic and clinical variables offers a novel approach for gaining insight into the effects of variables that are not explicitly contained in the training data. This approach has implications beyond multimodal explainability. Model developers could use it to better understand how aspects of their data are affecting the patterns learned by their models. Additionally, enhanced explainability of model decision-making could increase physicians’ and other relevant decision-makers’ trust of deep learning-based systems and subsequently increase the likelihood of the adoption of their recommendations. Scientists could also use the approach to help develop hypotheses for novel biomarkers. For example, if age were related to the explanations for a class and variable or the presence of an Alzheimer’s disease-related gene were related to the explanations for a brain region, researchers could later examine that relationship with future studies.

Our global multimodal explainability approach is well suited to multimodal electrophysiology data, as its key difference from standard ablation approaches is that it replaces modalities with noise like the artifact commonly found in electrophysiology data. Like our global ablation approach, our local ablation approach would, in its current state, only be applicable to electrophysiology data. However, our examination of the effect upon the output activation following ablation could easily be adapted to other domains. Our use of a gradient-based approach is more methodologically sound than the use of ablation methods, so applying it would likely be better than using ablation across many problems. Additionally, the ablation and gradient-based methods would each be better suited to different models. Unlike gradient methods, ablation is applicable to all deep learning classification frameworks. For example, our ablation methods would likely be more effective for long short-term memory networks than our LRP approach, while our LRP-based approach would likely be more effective for many CNN or multilayer perceptron architectures (33). While that is generally the case, implementing LRP or similar methods with some CNN architectures would be difficult, while the implementation of ablation would be relatively straightforward.

### Classification Performance

Our classifier performed well overall but somewhat below state-of-the-art classifiers (47). The classifier performed worst on NREM1. This makes sense given that NREM1 is the smallest class and that it can be similar to Awake and REM (15)(21). Additionally, NREM1 classification has normally been relatively poor in previous studies (20,21,27,40), so some studies have specifically developed methods that improve NREM1 classification (20). Although the Awake and NREM1 had similar numbers of samples, the classifier performed markedly better on Awake. Given that Awake EEG and EOG have features that are very different from those of NREM and that Awake EMG is different from REM EMG (15), it makes sense that the classifier would have an easier time learning to the classify Awake samples. Similar to previous studies (47)(27), the precision and F1 score of the classifier, but not the recall, was highest for NREM2.

### Global Results

Across methods, EEG was most important for identifying Awake, NREM2, NREM3, and REM. In contrast, EOG played a greater role in the correct classification of NREM1 samples. EMG was not very important to the classification of any stage. This result is not atypical, as previous studies have shown that using both EEG and EMG does not significantly improve classification performance for Awake, NREM2, and NREM3 relative to just using EEG (48). While global explanations for correct classification groups were similar across methods, explanations for incorrect groups tended to differ across methods.

### Subject-level Local Ablation and LRP Results over Time

The two local approaches had similar results for the 2-hour period of explanations that we output. In contrast to the global explanations, EOG was particularly important during Awake periods. This suggests that subject or subgroup-specific patterns of EOG activity exist within the Awake class that are obscured by global methods. It also supports existing findings that showed how EEG alone did not discriminate between Awake, NREM1, and REM as effectively as EEG with EOG and EMG (49). Additionally, previous studies have found that EOG is particularly important for identifying Awake (50) and can yield comparable classification performance to EEG (51). In contrast, EEG was important for discriminating NREM and REM samples. The importance of EEG for NREM and REM makes sense given that EEG patterns for NREM and REM differ greatly (15). However, it is interesting that this subject had higher Awake EOG than EEG importance. Globally, EEG tended to be more important for Awake than EOG. Moreover, visualizing the results over time enabled us to obtain higher resolution insight into the classifier than visualizing them globally. For example, they showed that EMG importance for the subject typically spiked when samples were incorrectly classified, which suggests that EMG adversely affected model performance.

### Statistical Analysis of Effects of Clinical and Demographic Variables upon Local Explanations

We assume that the relationships that are shared by both local explainability methods with demographic and clinical variables are more likely to be significant relationships than those identified by only one explainability method. As such, we only discuss the implications of those replicable patterns within the context of established research. Interestingly, subject sex has relationships with more modality correct classification group pairs than either medication or age, which could indicate that subject sex had stronger effects on the patterns learned by the classifier than the other variables. Given that males and females are highly effect could be attributed to the imbalance of male to female subjects present in the dataset.

Subject sex seemingly affected the NREM1 EEG patterns learned by the classifier. This reflects well-characterized aspects of sleep science. Namely, that adult women can have greater slow-wave EEG activity in NREM sleep stages than men (52)(53) and that, in general, there are differences in the EEG activity of men and women (54)(55). While sex was associated with the correct classification of NREM1, sex may also have adversely affected the EEG patterns learned by the classifier for several sleep stages, like Awake, NREM1, NREM2, and REM. Whereas the effects of sex on EEG was more associated with the incorrect classification of samples, both explainability methods indicated that sex likely affected the EOG patterns learned by the classifier for the correct classification of Awake, NREM2, NREM3, and REM. This highlights the possibility of EOG sex differences across most sleep stages. We have been unable to find many studies on the effects of sex upon EOG in sleep, so our results could prompt in-depth future studies on this topic. Both methods indicated that sex seemed to affect the EMG patterns learned for incorrectly classified samples. Medication seemed to affect the EEG of Awake, NREM2, and NREM3 samples similarly in both explainability methods. Previous studies have shown that benzodiazepines like temazepam (56) and other medications like duloxetine and desipramine (57) can have significant effects on EEG sleep stages and that temazepam, in particular, can have significant effects upon REM (58). Other studies have shown similar effects in monkey EEG (59). Our results further showed that medication significantly affected the patterns learned for REM EOG. Interestingly, medication may have been slightly related to the learning of EMG patterns that contributed to incorrect NREM classification. That there were inconsistent effects of medication upon EMG could fit with previous studies that purportedly analyzed EMG sleep data in monkeys but did not report any effects of medication (59). The effects of age on sleep are well characterized (52,53,60–62). In our study, age seemed to affect the EEG patterns learned for REM, similar to (63). However, age was also related to the learning of EEG patterns for multiple incorrect classification groups. This suggests that the model did not fully learn to address the underlying effects of age upon EEG across sleep stages. Interestingly, age had inconsistent effects upon the EOG patterns learned by the classifier. Age seemed to affect EMG patterns for NREM2 and NREM3.

### Limitations and Next Steps

The dataset that we used contained many subjects. Future studies might perform analyses comparing differences in importance across subjects, which could enable the identification of personalized sleep stage biomarkers (43). In this study, we used a simple CNN classifier, which made the implementation of LRP straightforward. However, using a simple CNN classifier also contributed to classification performance that was high but below the state of the art. Future studies with advanced classifiers might use the analyses that we employed to assist with the discovery of biomarkers and formulation of novel hypotheses related to sleep and other domains. Our classifier was originally developed for EEG sleep stage classification. As such, the architecture may not be optimized for extracting EOG and EMG features. This does not, in any way, adversely affect the quality of our explainability results. However, it prevents generalizable claims regarding the importance of one modality over another for the identification of sleep stages. Additionally, other GBFA methods could potentially replace LRP for multimodal explainability. Quality metrics like those presented in (64) could help rate the quality of each method for explaining classifiers trained on multimodal electrophysiology data. Additionally, previous studies have shown that applying multiple relevance rules sequentially to different parts of a network can yield less noisy explanations (65), and future studies might enhance explanation quality using that approach.

### Concluding Thoughts

In this study, we propose sleep stage classification as a testbed for developing novel multimodal explainability methods. Further, after training a classifier for multimodal sleep stage classification, we present a series of novel explainability methods for deep learning classifiers trained on multimodal time-series data. Our global ablation method is more domain-friendly than previous approaches that have replaced modalities with zeros. Our local ablation approach is, to the best of our knowledge, the first local multimodal ablation method to be developed, and our GBFA approach offers an alternative to ablation that has not previously been used for understanding modality importance. We show how the methods can be used to obtain insight into the importance of modalities to both incorrect and correct classification of samples. We find that EEG was most important to the identification of most sleep stages while EOG is most important of the identification of NREM1. We show how local methods can help identify differences in subject-level explanations that differ from global explanations and that could potentially be used to identify personalized biomarkers in future studies. For example, one subject showed higher levels of EOG importance in Awake samples than EEG importance. Importantly, we also developed a novel analysis approach and found that subject sex had more significant relationships with patterns learned by the classifier for each sleep stage relative to other clinical and demographic variables. More broadly, the approach could help illuminate the effects of those variables upon different classes (e.g., sleep stages or disease conditions). Our study greatly enhances the degree of insight that can be obtained from the typically black-box models of the growing field of multimodal classification.

## Methods

In this section, we describe our data, preprocessing, model architecture and training approach, and explainability methods.

### Description of Data

We utilized Sleep Telemetry data from the Sleep-EDF Expanded Database (16) on Physionet (66). The database can be downloaded at (67) and has been used in previous sleep stage classification studies (5,19,25,29). The dataset has 44 approximately 9-hour recordings from 22 subjects (15 female and 7 male) with primary sleep onset insomnia (68). Subject age had a mean of 40.18 years and a standard deviation of 18.09 years. Figure 8 shows subject demographics. All subjects had two recordings – one following the administration of a placebo and one following temazepam administration. Temazepam belongs to a class of drugs called benzodiazepines which amplify the effects of the neurotransmitter y-aminobutyric acid (GABA). GABA is the most common neurotransmitter in the central nervous system. It is inhibitory in nature and produces a calming effect on the brain (69). It is commonly used to treat insomnia and affects electrophysiology activity. Each recording contained data from 2 EEG electrodes, 1 EOG electrode, and 1 EMG electrode. Data was recorded at a 100 Hertz (Hz) sampling frequency. The two EEG electrodes were FPz-Cz and Pz-Oz, according to (70). However, like previous studies (5,20,25,29,71), we used only the Fpz-Cz for our analysis. A 1-Hz marker was derived from the recording device that indicated the presence of recording errors. Using the Rechtschaffen and Kales standard (72), expert technicians assigned 30-second data epochs to one of seven categories: Movement, Awake, REM, NREM1, NREM2, NREM3, and NREM4. In our study, we merged NREM3 and NREM4 into a single NREM3 class (15). We removed all movement samples and samples containing recording errors.

**Figure 8.**
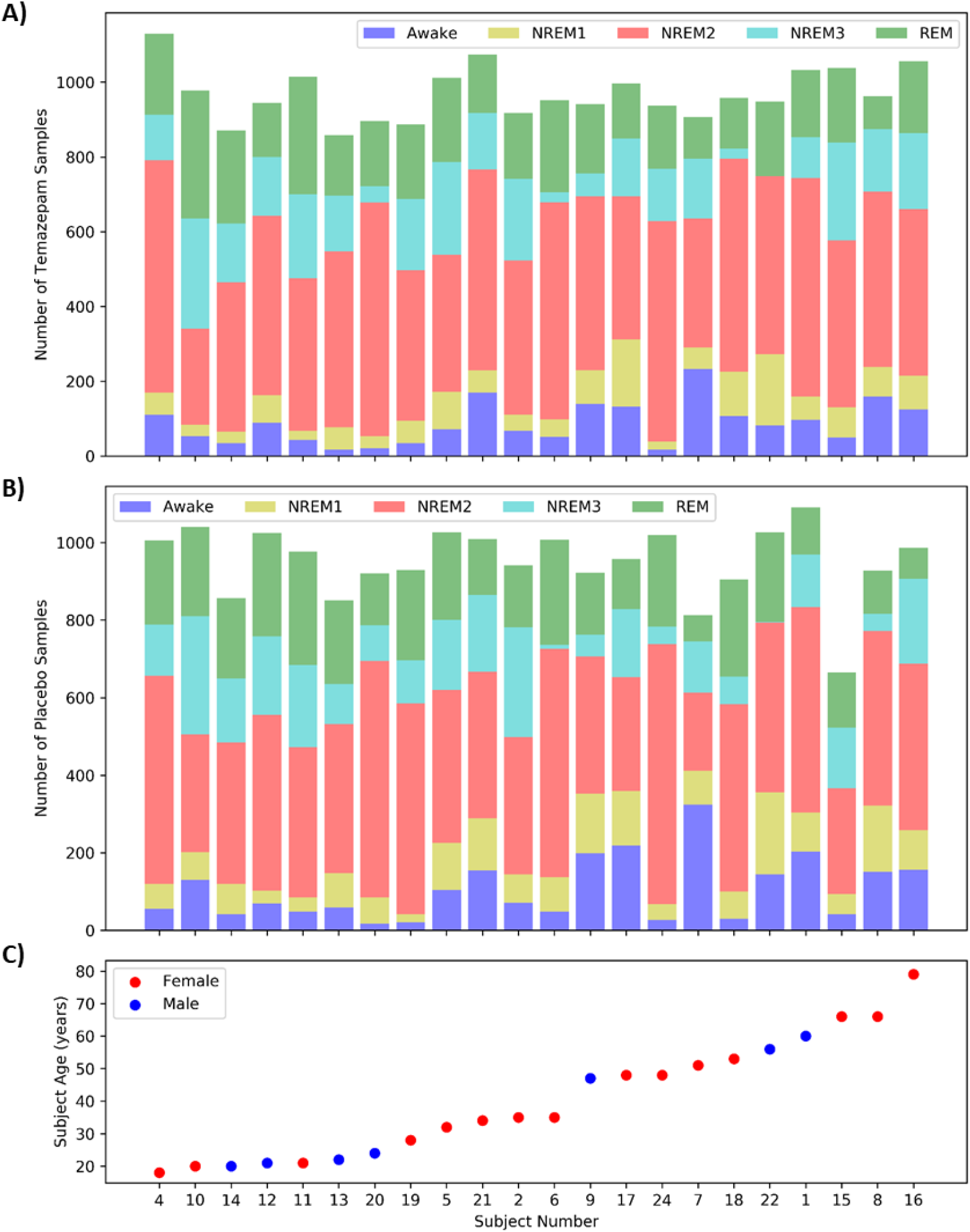
Distribution of Samples and Subject Demographics. Panels A and B show the distributions of temazepam and placebo samples, respectively, for each subject. Panel C shows the age and sex of each subject, with the subjects arranged from youngest to oldest. Each panel shares the same x-axis.

### Description of Data Preprocessing

Based on the data annotation, we segmented the data into 30-second samples. Within each recording, we z-scored each electrode individually to improve the identification of patterns across subjects. After segmentation and sample removal, our dataset had 42,218 samples. The dataset was highly imbalanced with Awake, NREM1, NREM2, NREM3, and REM classes having 9.97%, 8.53%, 46.8%, 14.92%, and 19.78% of the dataset, respectively. Figure 8 shows the distribution of samples in each class for each subject.

### Description of 1D-CNN

#### Model Architecture and Training

We adapted a CNN architecture initially developed for EEG classification (73). Details on the architecture are shown in Figure 9. We implemented the architecture in Keras (74) with a TensorFlow (75) backend. We used 10-fold cross-validation with 17, 2, and 3 subjects being randomly assigned to training, validation, and test sets in each fold. We used categorical cross entropy loss and weighted each class to account for class imbalances when training the classifier. We used a batch size of 100 with shuffling after each epoch. We used the Adam optimizer (76) with an adaptive learning rate. Starting with an initial learning rate of 0.001, the optimizer decreased its step size by a factor of 10 after every 5 epochs in which validation accuracy did not improve. We used early stopping to end training after validation accuracy plateaued for 20 epochs with a maximum of 100 epochs and used model checkpoints to select the model from each fold that obtained the highest validation accuracy. We used the selected models for testing and explainability.

**Figure 9.**
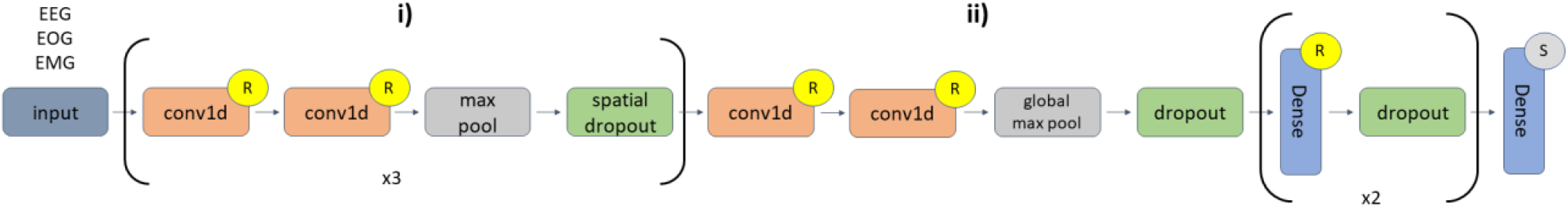
CNN Architecture. Layers **i)** of the diagram repeat 3 times. In **i)** there are 6 1D-convolutional (conv1d) layers in total. The first two conv1d layers (number of filters = 16, kernel size = 5) are followed by max pooling (pool size = 2) and spatial dropout (rate = 0.01). The second two conv1d layers (number of filters = 32, kernel size = 3) are followed by max pooling (pool size = 2) and a spatial dropout (rate = 0.01). The third pair of conv1d layers (number of filters = 32, kernel size = 3) are followed by max pooling (pool size = 2) and spatial dropout (rate = 0.01). In **ii)**, the last two conv1d layers (number of filters = 256, kernel size = 3) are followed by global max pooling and dropout (rate = 0.01). The first two dense layers (number of nodes = 64) have dropout rates of 0.1 and 0.05, respectively. The last dense layer has 5 nodes. An “R” or an “S” indicates that a layer is followed by ReLU or Softmax activation functions, respectively.

#### Model Performance Evaluation

When evaluating model test performance, we sought to account for class imbalances. We generated a confusion matrix and calculated precision (Equation 1), recall (Equation 2), and the F1 score (Equation 3) for each class. We calculated the mean and standard deviation of the metrics across folds.

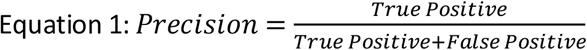

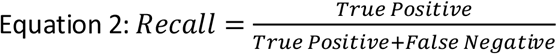

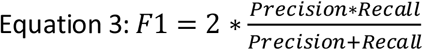

### Description of Global Ablation Approaches

In this study, we applied two global ablation approaches for insight into the relative importance of each modality to each class. We presented a novel approach for global ablation that is unique to the electrophysiology domain and that we originally presented in (12). We compared our novel approach to an ablation approach that has been used in previous multimodal classification studies (2)(41).

#### Standard Global Ablation Approach

The typical ablation approach that we applied followed several key steps. (1) We calculated a confusion matrix within a single test fold. (2) We replaced all values for a modality with zeros. (3) We calculated a confusion matrix for the classifier on the modified data. (4) We calculated the percent change in samples assigned to each classification group (Equation 4). (5) We repeated steps 2 through 4 for each modality. (6) We repeated steps 1 through 5 for each test fold.

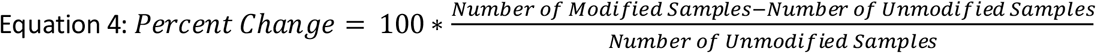

#### Our Novel Global Ablation Approach

In this study, we propose a novel ablation approach for multimodal electrophysiology analysis that involves replacing modalities in a manner that mimics line-related noise. In Step 2 of the previously detailed ablation process, we replace a modality with a combination of a sinusoid and Gaussian noise. We use a sinusoid with a frequency of 40 Hz because 60 Hz noise at a sampling rate of 100 Hz would alias and appear at 40 Hz. The sinusoid had an amplitude of 0.1, and the Gaussian noise was generated from a distribution with a mean of 0 and standard deviation of 0.1.

#### Statistical Comparison

To determine whether our novel line-related noise ablation yielded results significantly different from standard ablation, we performed a series of two-tailed t-tests. Within each modality, we compared the ablation importance values in each classification group for each method across folds.

### Description of Novel Local Ablation Approach

We applied a novel local ablation approach for insight into the modality importance to the classification of each sample. We originally presented the approach in (13). Our novel ablation approach is similar to the global approach described in the previous section. (1) We obtained the top-class probability for a particular sample. (2) We modified the sample by ablating a modality. (3) We obtained the classification probability of the modified sample for the previously identified top class. (4) We computed the percent change in classification probability (Equation 5). (5) We repeated steps 2 through 4 for each modality. (6) We repeated steps 2 through 5 for each sample in a fold. (7) We repeated steps 2 through 6 for each fold.

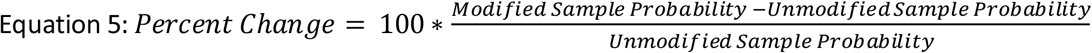

Because there were no preexisting local approaches for comparison, we examined how comparable our local ablation results were to the previously described global methods. In addition to generating local visualizations of our local ablation results, we formed a global estimate by calculating the mean absolute percent change in classification probability for each fold, modality, and classification group.

### Description of Layer-Wise Relevance Propagation Analysis

Layer-wise relevance propagation (LRP) (31) is a popular GBFA method (46) originally developed for image analysis. Since its initial development, LRP has been used in raw electrophysiology analysis (77) and other neuroscience domains (33,78,79). Other GBFA methods have also been applied to multimodal time-series classification (43), and we previously applied LRP to multimodal electrophysiology analysis (11). Local ablation methods, like sensitivity analysis (28) and many GBFA methods, show what features or time points make a sample more or less like the patterns learned by the classifier for a particular class. LRP shows what features or time points are actually used by the classifier for its classification and indicates their importance (80). We implemented LRP using the Innvestigate library (81).

LRP is a local explainability method but has also been used for global importance estimates (11,33). LRP involves the following steps. (1) A sample is passed through a network and assigned a particular class. (2) A total relevance of 1 is placed at the output node of the assigned class. (3) The total relevance is propagated through the network using relevance rules until the relevance is distributed across the input features. A key characteristic of LRP is that the total relevance value of 1 is conserved as it is propagated through the network, so the relevance assigned to the input features should sum to approximately 1.

There can be both negative and positive relevance depending upon the relevance rule that is used. Negative relevance indicates the features of an input sample that support a sample being classified as a class other than that which it was assigned. For example, if the value of feature *x* for a sample classified as class 1 were more in alignment with class 2, that feature would receive negative relevance. Positive relevance indicates the features of an input sample that support a sample being classified as its assigned class. For instance, if a sample belongs to class 1 and a particular pattern in that sample provides evidence for class 1, that pattern would be assigned positive relevance. Equation 6 shows the basic LRP rule:

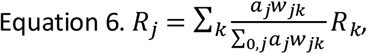

where *k* indicates a node that is one of *k* nodes in a layer deeper in a network and *j* indicates a node in the layer to which relevance is being propagated. *R*_*k*_ indicates the total relevance assigned to a node in a deeper layer, and *R*_*j*_ indicates the total relevance that will be assigned to a node in a shallower layer. The variables *a*_*j*_ and *w*_*jk*_ indicate the activation output of the layer j and the value of the weight connecting the node in layer *j* and node in layer *k*. The numerator indicates a portion of the effect that the node in layer *j* has upon the node in layer *k*, and the denominator indicates the total effect of all nodes in layer *j* upon the node in layer *k*. This combined with the summation Σ_k_ indicates that the relevance assigned to the node in layer *j* is the sum of the fraction of the effect of the node in layer *j* upon all of the nodes in layer *k* multiplied by their respective relevance.

The basic LRP rule allows for both positive and negative relevance to be propagated through the network and often yields very noisy explanations. In our study, we used the ε-rule and αβ-rule. As shown in Equation 7, the ε-rule is identical to the basic rule except for the added term ε.

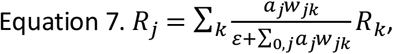

where ε enables relevance to be filtered when propagated through the network. A larger ε shrinks the amount of relevance propagated backwards for nodes that would otherwise be assigned low relevance. In effect, this reduces the noisiness of the explanations. We used the ε-rule with an ε of 0.01 and 100.

We also used the αβ-rule shown in Equation 8,

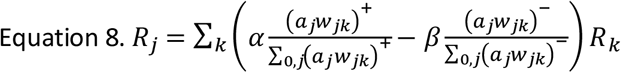

where the relevance is split into positive and negative portions when propagated backwards. The variables α and β control how much positive and negative relevance are propagated backwards, respectively. In our study, we used the rule with α = 1 and β = 0, to only propagate positive relevance.

While conducting our LRP analysis, we generated a global estimation of importance by calculating the percent of absolute relevance assigned to each modality. We computed this value for each classification group in each fold. We also visualized how the percent of relevance varied over time.

### Description of Additional Statistical Analyses

We performed a series of statistical analyses with the local ablation and LRP (ε-rule with ε = 100) explanations for insight into the effects of demographic and clinical variables upon the classifier. We trained an ordinary least squares regression model with age, medication, and sex as the independent variables and with the absolute importance (percent change in activation for local ablation and relevance for LRP) for a modality and classification group as the dependent variable. This enabled us to control for interaction effects. For LRP, we used the percent of absolute relevance assigned to each modality for each sample. After training the model, we obtained the resulting coefficients and p-values for each class. The sign of the coefficients identified the direction of the importance difference. After obtaining p-values, we performed false discovery rate (FDR) correction (α = 0.05) with the 25 p-values (i.e., 5 classes x 5 classes) associated with each clinical or demographic variable to account for multiple comparisons.

## Acknowledgements

This work was funded by NIH grant R01EB006841.

## Supplementary Material

**Supplementary Figure 1.**
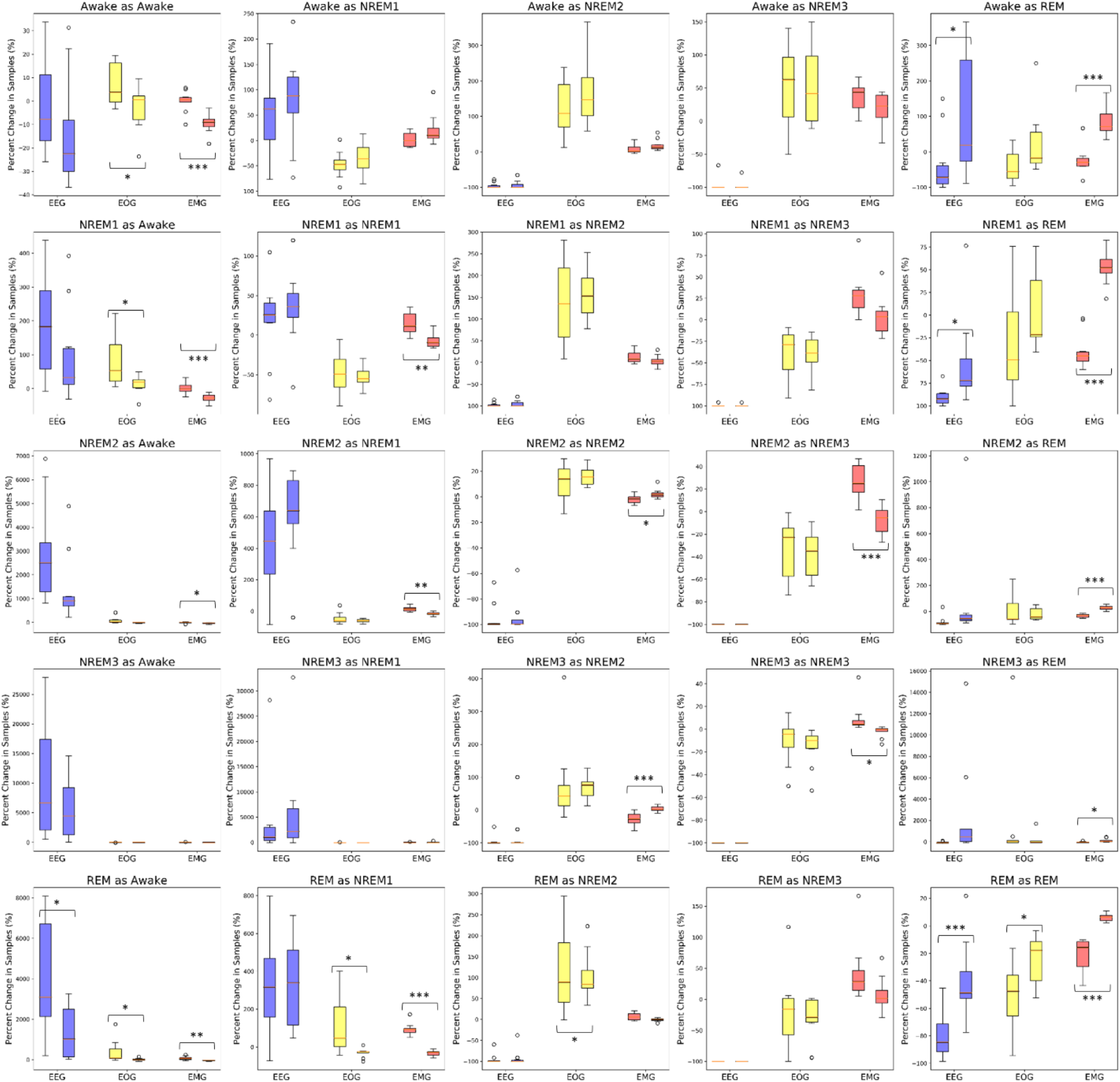
Global Ablation Explainability Results for All Classification Groups. The above boxplots show the percent change in samples assigned to each classification group following perturbation for each fold. Blue, yellow, and red boxes indicate EEG, EOG, and EMG perturbation, respectively. The leftmost box of each pair shows the results for our noise-related ablation method. The rightmost box is for the typical zero-out ablation approach. Some pairs of boxes are labeled with *, **, or ***, which correspond to a significance value of p < 0.05, p < 0.01, and p < 0.001, respectively.

**Supplementary Figure 2.**
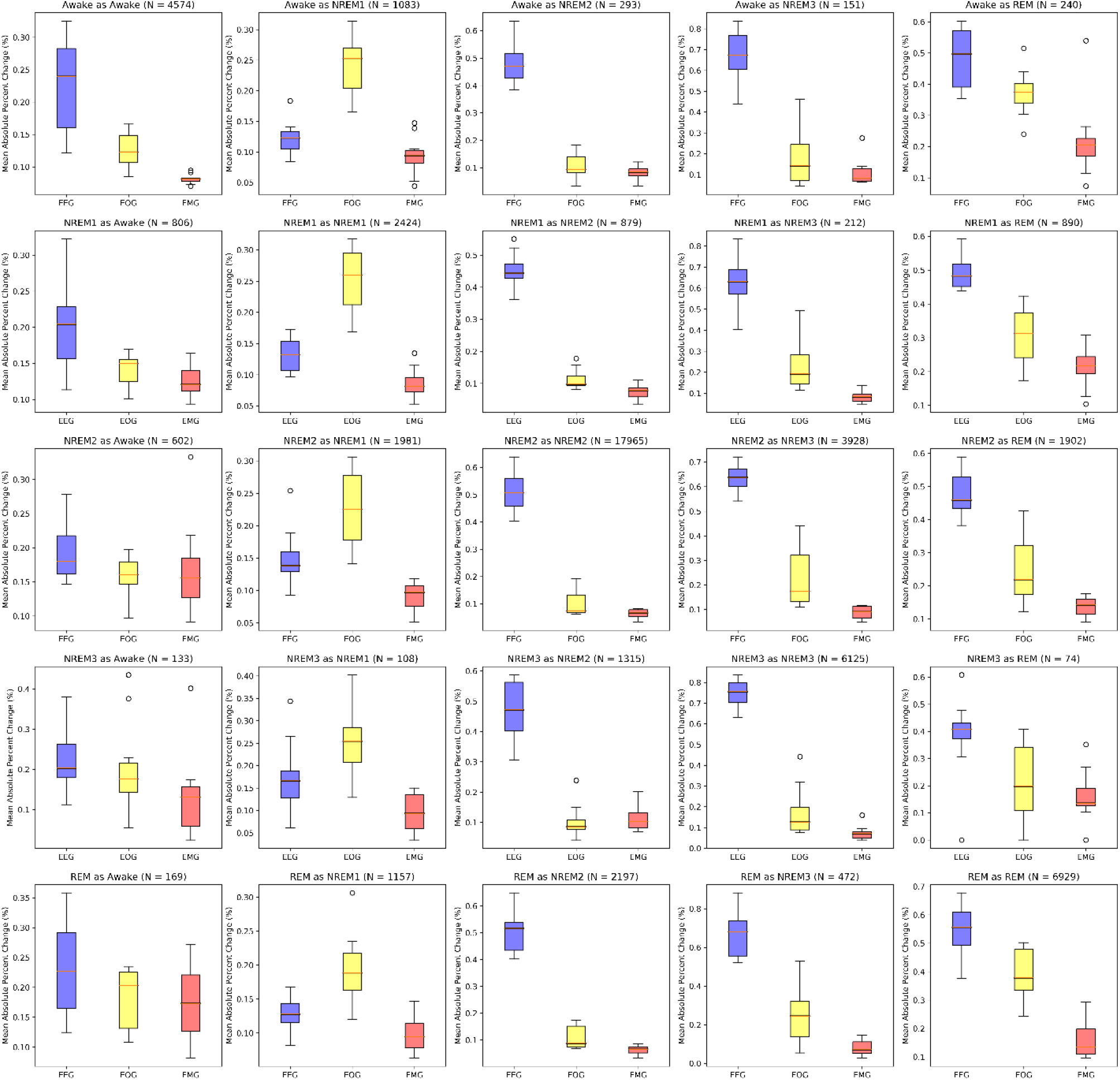
Local Ablation Results Showing Global Estimation of Modality Importance for All Classification Groups. The above boxplots show the percent change in activation following perturbation for samples in each classification group across folds. Blue, yellow, and red boxes indicate EEG, EOG, and EMG perturbation, respectively.

**Supplementary Figure 3.**
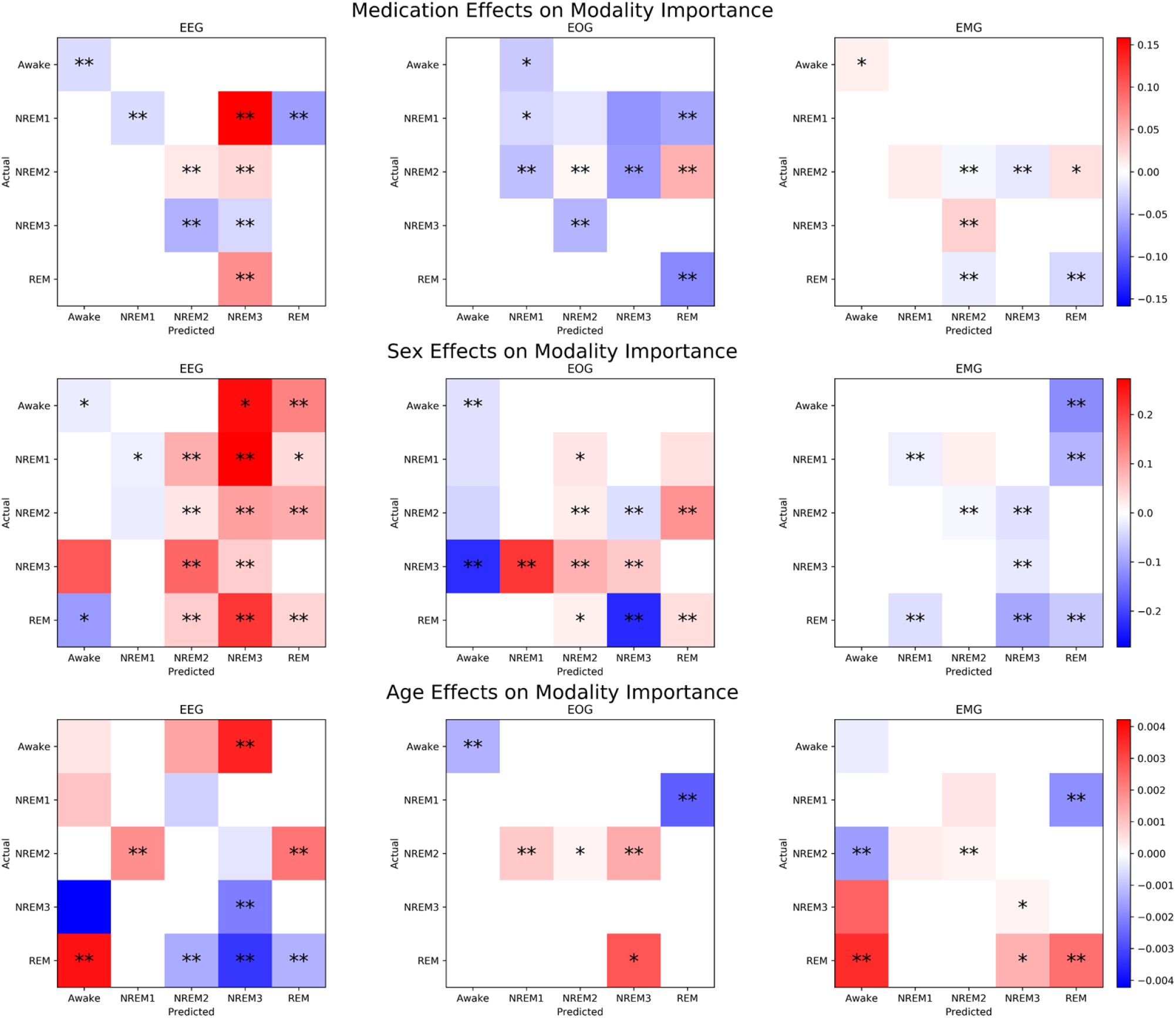
Effects of Clinical and Demographic Variables upon Local Ablation Modality Importance for All Classification Groups. The top, middle, and bottom rows of panels indicate the effects of medication, sex, and age, respectively, upon the explanations for each modality and classification group. The x-axis of each panel indicates the predicted class, and the y-axis indicates the actual class. The heatmaps indicate the size of the coefficient values resulting from the regression analysis. White squares have insignificant p-values. Squares with color have significant p-values (p<0.05), and squares that have one or two asterisks have p-values of p < 0.01 or p < 0.001, respectively. For medication, a positive coefficient value indicates that temazepam samples had more importance than placebo samples. For subject sex, a positive coefficient value indicates that female samples had more importance than male samples, and for age, a positive coefficient indicates that importance increased with age.

**Supplementary Figure 4.**
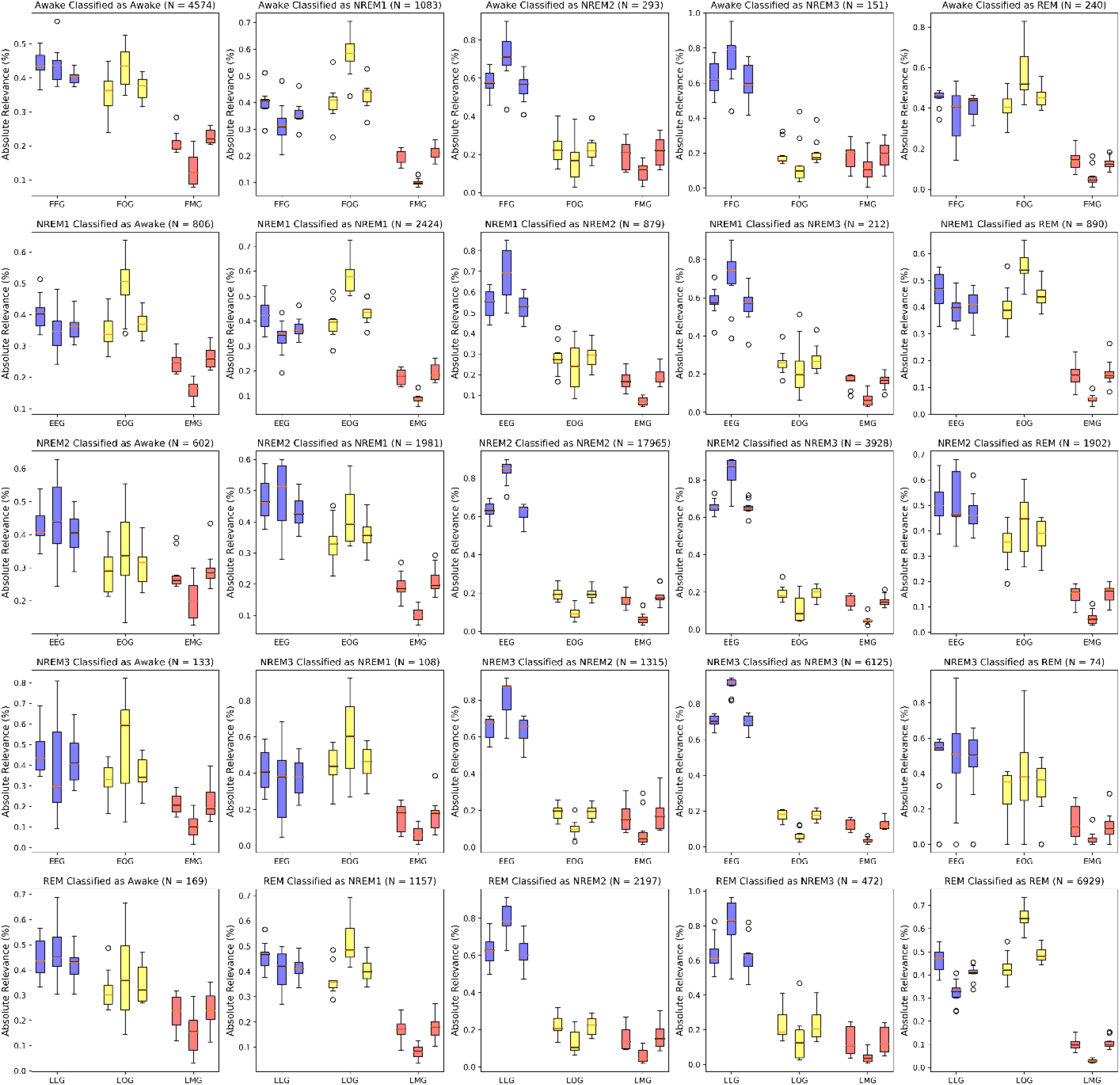
LRP-based Global Estimation of Modality Importance for All Classification Groups. The boxplots show the LRP explainability results for each classification group across folds. Blue, yellow, and red boxes show the importance of EEG, EOG, and EMG, respectively. From left to right, within each group of three boxes are the relevance results for the LRP ε-rule (0.01), ε-rule (100), and αβ-rule. The number of samples in each classification group is included in the title of each panel. The panels on the left-to-right diagonal show correct classification groups, and off-diagonal panels indicate incorrect classification groups.

**Supplementary Figure 5.**
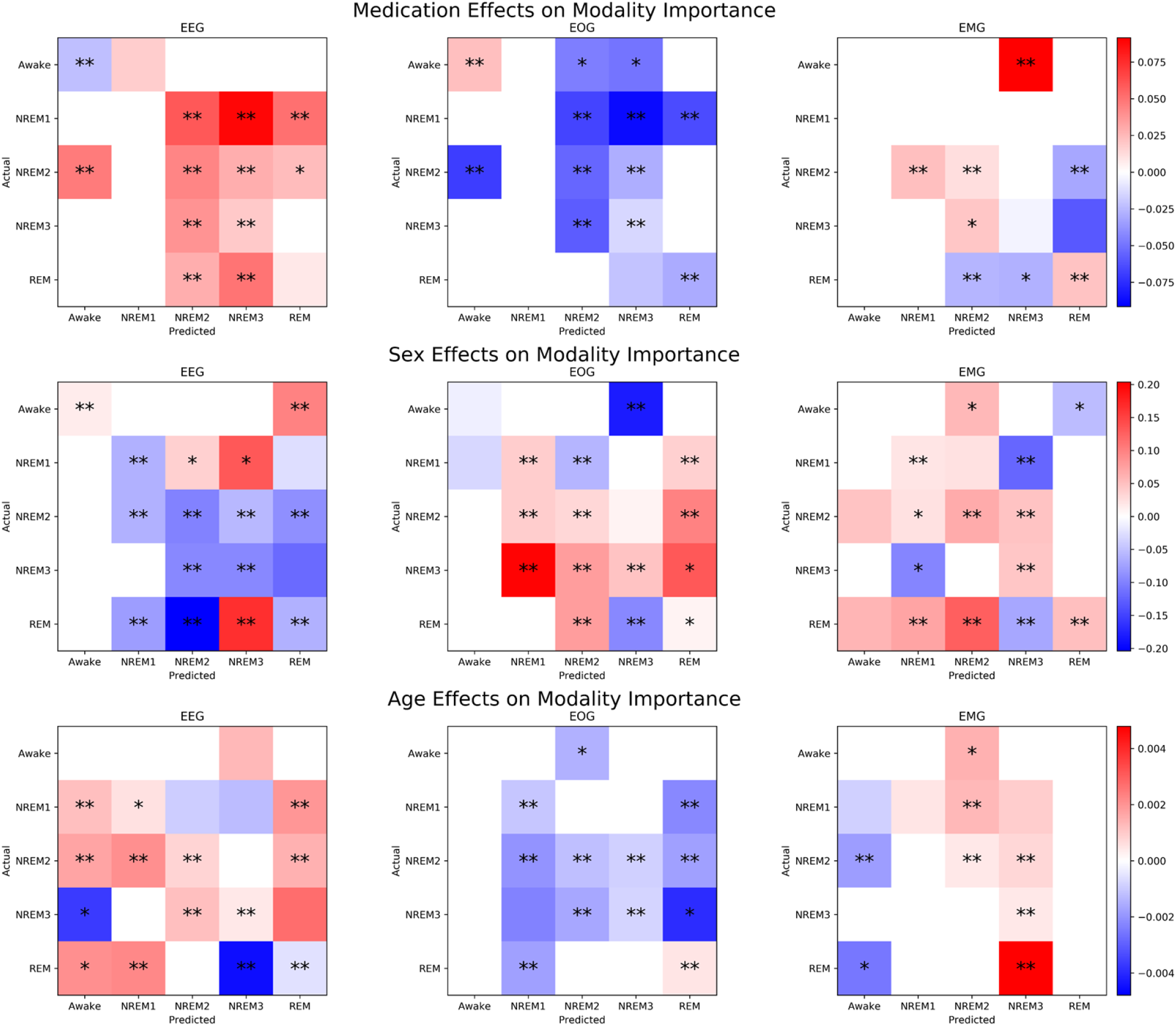
Effects of Clinical and Demographic Variables upon LRP Relevance for All Classification Groups. The top, middle, and bottom rows of panels indicate the effects of medication, sex, and age, respectively, upon the explanations for each modality and classification group. The x -axis of each panel indicates the predicted class, and the y-axis indicates the actual class. The heatmaps indicate the size of the coefficient values resulting from the regression analysis. White squares have insignificant p-values. Squares with color have significant p-values (p<0.05), and squares that have one or two asterisks have p-values of p < 0.01 or p < 0.001, respectively. For medication, a positive coefficient value indicates that temazepam samples had more relevance than placebo samples. For subject sex, a positive coefficient value indicates that female samples had more relevance than male samples, and for age, a positive coefficient indicates that relevance increased with age.

## References

1. Zhai B, Perez-Pozuelo I, Clifton EAD, Palotti J, Guan Y. Making Sense of Sleep: Multimodal Sleep Stage Classification in a Large, Diverse Population Using Movement and Cardiac Sensing. Proc ACM Interactive, Mobile, Wearable Ubiquitous Technol. 2020;4(2).

2. Lin J, Pan S, Lee CS, Oviatt S. An Explainable Deep Fusion Network for Affect Recognition Using Physiological Signals. In: Proceedings of the 28th ACM International Conference on Information and Knowledge Management. 2019. p. 2069–72.

3. Mellem MS, Liu Y, Gonzalez H, Kollada M, Martin WJ, Ahammad P. Machine Learning Models Identify Multimodal Measurements Highly Predictive of Transdiagnostic Symptom Severity for Mood, Anhedonia, and Anxiety. Biol Psychiatry Cogn Neurosci Neuroimaging. 2020 Jan 1;5(1):56–67.

4. Wang IN, Lee CH, Kim HJ, Kim H, Kim DJ. An Ensemble Deep Learning Approach for Sleep Stage Classification via Single-channel EEG and EOG. Int Conf ICT Converg. 2020;2020-Octob:394–8.

5. Phan H, Andreotti F, Cooray N, Chen OY, De Vos M. Joint Classification and Prediction CNN Framework for Automatic Sleep Stage Classification. IEEE Trans Biomed Eng. 2019;66(5):1285–96.

6. Kwon YH, Shin SB, Kim SD. Electroencephalography based fusion two-dimensional (2D)-convolution neural networks (CNN) model for emotion recognition system. Sensors (Switzerland). 2018 May 1;18(5).

7. Niroshana SMI, Zhu X, Chen Y, Chen W. Sleep Stage Classification Based on EEG, EOG, and CNN - GRU Deep Learning Model. 2019 IEEE 10th Int Conf Aware Sci Technol iCAST 2019 - Proc. 2019;1–7.

8. Li Y, Yang X, Zhi X, Zhang Y, Cao Z. Automatic Sleep Stage Classification Based on Two-channel EOG and One-channel EMG. Res Sq [Internet]. :1–15. Available from: https://www.researchsquare.com/article/rs-491468/latest?utm_source=researcher_app&utm_medium=referral&utm_campaign=RESR_MRKT_Researcher_inbound

9. Zhang D, Wang Y, Zhou L, Yuan H, Shen D. Multimodal classification of Alzheimer’s disease and mild cognitive impairment. Neuroimage. 2011 Apr 1;55(3):856–67.

10. Sullivan HR, Schweikart SJ. Are current tort liability doctrines adequate for addressing injury caused by AI? AMA J Ethics. 2019;21(2):160–6.

11. Ellis CA, Carbajal DA, Zhang R, Miller RL, Calhoun VD, Wang MD. An Explainable Deep Learning Approach for Multimodal Electrophysiology Classification. bioRxiv. 2021;12–5.

12. Ellis CA, Zhang R, Carbajal DA, Miller RL, Calhoun VD, Wang MD. Explainable Sleep Stage Classification with Multimodal Electrophysiology Time-series. bioRxiv. 2021;0–3.

13. Ellis CA, Carbajal DA, Zhang R, Sendi MSE, Miller RL, Calhoun VD, et al. A Novel Local Ablation Approach For Explaining Multimodal Classifiers. bioRxiv. 2021;1–6.

14. Ellis CA, Miller RL, Calhoun VD, Wang MD. A Gradient-based Approach for Explaining Multimodal Deep Learning Classifiers. In: 2021 IEEE 21st International Conference on Bioinformatics and Bioengineering (BIBE). IEEE; 2021. p. 0–5.

15. Iber C, Ancoli-Israel S, Chesson AL, Quan SF. The AASM Manual for Scoring of Sleep and Associated Events: Rules, Terminology, and Technical Specifications. 2007.

16. Kemp B, Zwinderman AH, Tuk B, Kamphuisen HAC, Oberye JJL. Analysis of a sleep-dependent neuronal feedback loop: the slow-wave microcontinuity of the EEG. IEEE Trans Biomed Eng. 2000;47(9):1185–94.

17. Khalighi S, Sousa T, Santos JM, Nunes U. ISRUC-Sleep: A comprehensive public dataset for sleep researchers. Comput Methods Programs Biomed [Internet]. 2016;124(October 2017):180–92. Available from: http://dx.doi.org/10.1016/j.cmpb.2015.10.013

18. Quan SF, Howard B V., Iber C, Kiley JP, Nieto FJ, O’Connor GT, et al. The Sleep Heart Health Study: Design, rationale, and methods. Sleep. 1997;20(12):1077–85.

19. Rahman MM, Bhuiyan MIH, Hassan AR. Sleep stage classification using single-channel EOG. Comput Biol Med [Internet]. 2018;102(June):211–20. Available from: https://doi.org/10.1016/j.compbiomed.2018.08.022

20. Michielli N, Acharya UR, Molinari F. Cascaded LSTM recurrent neural network for automated sleep stage classification using single-channel EEG signals. Comput Biol Med [Internet]. 2019;106(December 2018):71–81. Available from: https://doi.org/10.1016/j.compbiomed.2019.01.013

21. Tsinalis O, Matthews PM, Guo Y. Automatic Sleep Stage Scoring Using Time-Frequency Analysis and Stacked Sparse Autoencoders. Ann Biomed Eng. 2016;44(5):1587–97.

22. Rojas I, Joya G, Catala A. Deep Learning Using EEG Data in Time and Frequency Domains for Sleep Stage Classification. In: International Conference on Neural Information Processing. 2017. p. V–VII.

23. Aboalayon KAI, Almuhammadi WS, Faezipour M. A comparison of different machine learning algorithms using single channel EEG signal for classifying human sleep stages. In: 2015 IEEE Long Island Systems, Applications and Technology Conference, LISAT 2015. IEEE; 2015. p. 1–6.

24. Sors A, Bonnet S, Mirek S, Vercueil L, Payen JF. A convolutional neural network for sleep stage scoring from raw single-channel EEG. Biomed Signal Process Control. 2018;42:107–14.

25. Mousavi S, Afghah F, Rajendra Acharya U. SleepEEGNet: Automated Sleep Stage Scoring with Sequence to Sequence Deep Learning Approach. arXiv. 2019;1–15.

26. Eldele E, Chen Z, Liu C, Wu M, Kwoh CK, Li X, et al. An Attention-Based Deep Learning Approach for Sleep Stage Classification with Single-Channel EEG. IEEE Trans Neural Syst Rehabil Eng. 2021;29:809–18.

27. Supratak A, Dong H, Wu C, Guo Y. DeepSleepNet: A model for automatic sleep stage scoring based on raw single-channel EEG. IEEE Trans Neural Syst Rehabil Eng. 2017;25(11):1998–2008.

28. Simonyan K, Vedaldi A, Zisserman A. Deep Inside Convolutional Networks: Visualising Image Classification Models and Saliency Maps. 2013 Dec 20; Available from: http://arxiv.org/abs/1312.6034

29. Vilamala A, Madsen KH, Hansen LK. Deep convolutional neural networks for interpretable analysis of EEG sleep stage scoring. IEEE Int Work Mach Learn Signal Process MLSP. 2017;2017-Septe(659860):1–6.

30. Ruffini G, Ibañez D, Castellano M, Dubreuil-Vall L, Soria-Frisch A, Postuma R, et al. Deep Learning With EEG Spectrograms in Rapid Eye Movement Behavior Disorder. Front Neurol. 2019 Jul 30;10.

31. Bach S, Binder A, Montavon G, Klauschen F, Müller KR, Samek W. On pixel-wise explanations for non-linear classifier decisions by layer-wise relevance propagation. PLoS One. 2015 Jul 10;10(7).

32. Chen Y, Gong C, Hao H, Guo Y, Xu S, Zhang Y, et al. Automatic Sleep Stage Classification Based on Subthalamic Local Field Potentials. IEEE Trans Neural Syst Rehabil Eng. 2019;27(2):118–28.

33. Ellis CA, Sendi MS, Willie JT, Mahmoudi B. Hierarchical Neural Network with Layer-wise Relevance Propagation for Interpretable Multiclass Neural State Classification. In: 10th International IEEE/EMBS Conference on Neural Engineering (NER). 2021. p. 18–21.

34. Ellis CA, Miller RL, Calhoun VD. A Novel Local Explainability Approach for Spectral Insight into Raw EEG-Based Deep Learning Classifiers. In: bioRxiv. 2021. p. 0–5.

35. Ellis CA, Miller RL, Calhoun VD. A Gradient-based Spectral Explainability Method for EEG Deep Learning Classifiers. In: bioRxiv. 2021. p. 1–6.

36. Ellis CA, Sendi MSE, Miller R, Calhoun V. A Novel Activation Maximization-based Approach for Insight into Electrophysiology Classifiers. In: bioRxiv. 2021.

37. Ellis CA, Miller RL, Calhoun VD. A Model Visualization-based Approach for Insight into Waveforms and Spectra Learned by CNNs. bioRxiv. 2021;1–4.

38. Barnes LD, Lee K, Kempa-Liehr AW, Hallum LE. Detection of sleep apnea from single-channel electroencephalogram (EEG) using an explainable convolutional neural network. bioRxiv. 2021;

39. Nahmias DO, Kontson KL. Easy Perturbation EEG Algorithm for Spectral Importance (easyPEASI): A Simple Method to Identify Important Spectral Features of EEG in Deep Learning Models. In: Proceedings of the 26th ACM SIGKDD International Conference on Knowledge Discovery & Data Mining [Internet]. New York, NY, USA: ACM; 2020. p. 2398–406. Available from: https://dl.acm.org/doi/10.1145/3394486.3403289

40. Chambon S, Galtier MN, Arnal PJ, Wainrib G, Gramfort A. A deep learning architecture for temporal sleep stage classification using multivariate and multimodal time series. IEEE Trans Neural Syst Rehabil Eng. 2018;26(4):758–69.

41. Pathak S, Lu C, Nagaraj SB, van Putten M, Seifert C. STQS: Interpretable multi-modal Spatial-Temporal-seQuential model for automatic Sleep scoring. Artif Intell Med [Internet]. 2021;114(January):102038. Available from: https://doi.org/10.1016/j.artmed.2021.102038

42. Lajnef T, Chaibi S, Ruby P, Aguera PE, Eichenlaub JB, Samet M, et al. Learning machines and sleeping brains: Automatic sleep stage classification using decision-tree multi-class support vector machines. J Neurosci Methods [Internet]. 2015;250:94–105. Available from: http://dx.doi.org/10.1016/j.jneumeth.2015.01.022

43. Porumb M, Stranges S, Pescapè A, Pecchia L. Precision Medicine and Artificial Intelligence: A Pilot Study on Deep Learning for Hypoglycemic Events Detection based on ECG. Sci Rep. 2020;10(1):1–16.

44. Selvaraju RR, Cogswell M, Das A, Vedantam R, Parikh D, Batra D. Grad-CAM: Visual Explanations from Deep Networks via Gradient-Based Localization. Int J Comput Vis. 2020;128(2):336–59.

45. Molnar C. Interpretable Machine Learning A Guide for Making Black Box Models Explainable [Internet]. 2018th-08–14th ed. Lean Pub; 2018. Available from: http://leanpub.com/interpretable-machine-learning

46. Ancona M, Ceolini E, Öztireli C, Gross M. Towards Better Understanding of Gradient-based Attribution Methods for Deep Neural Networks. In: International Conference on Learning Representations. 2018. p. 1–16.

47. Chambon S, Galtier MN, Arnal PJ, Wainrib G, Gramfort A. A deep learning architecture for temporal sleep stage classification using multivariate and multimodal time series. arXiv. 2017;26(4):758–69.

48. Kim H, Choi S. Automatic Sleep Stage Classification Using EEG and EMG Signal. Int Conf Ubiquitous Futur Networks, ICUFN. 2018;2018-July:207–12.

49. Estrada E, Nazeran H, Barragan J, Burk JR, Lucas EA, Behbehani K. EOG and EMG: Two important switches in automatic sleep stage classification. In: Annual International Conference of the IEEE Engineering in Medicine and Biology - Proceedings. IEEE; 2006. p. 2458–61.

50. Pettersson K, Müller K, Tietäväinen A, Gould K, Hæggström E. Saccadic eye movements estimate prolonged time awake. J Sleep Res. 2019;28(2):1–13.

51. Ganesan RA, Jain R. Binary State Prediction of Sleep or Wakefulness Using EEG and EOG Features. 2020 IEEE 17th India Counc Int Conf INDICON 2020. 2020;

52. Ehlers CL, Kupfer DJ. Slow-wave sleep: Do young adult men and women age differently? J Sleep Res. 1997;6(3):211–5.

53. Mourtazaev MS, Kemp B, Zwinderman AH, Kamphuisen HAC. Age and gender affect different characteristics of slow waves in the sleep EEG. Sleep. 1995;18(7):557–64.

54. Bučková B, Brunovský M, Bareš M, Hlinka J. Predicting Sex From EEG: Validity and Generalizability of Deep-Learning-Based Interpretable Classifier. Front Neurosci. 2020;14(October):1–7.

55. Armitage R, Hoffmann RF. Sleep EEG, depression and gender. Sleep Med Rev. 2001;5(3):237–46.

56. Bastien CH, LeBlanc M, Carrier J, Morin CM. Sleep EEG power spectra, insomnia, and chronic use of benzodiazepines. Sleep. 2003;26(3):313–7.

57. Chalon S, Pereira A, Lainey E, Vandenhende F, Watkin JG, Staner L, et al. Comparative effects of duloxetine and desipramine on sleep EEG in healthy subjects. Psychopharmacology (Berl). 2005;177(4):357–65.

58. Pagel JF, Farnes BL. Medications for the treatment of sleep disorders: An overview. Prim Care Companion J Clin Psychiatry. 2001;3(3):118–25.

59. Authier S, Bassett L, Pouliot M, Rachalski A, Troncy E, Paquette D, et al. Effects of amphetamine, diazepam and caffeine on polysomnography (EEG, EMG, EOG)-derived variables measured using telemetry in Cynomolgus monkeys. J Pharmacol Toxicol Methods [Internet]. 2014;70(1):86–93. Available from: http://dx.doi.org/10.1016/j.vascn.2014.05.003

60. Chinoy ED, Frey DJ, Kaslovsky DN, Meyer FG, Wright KP. Age-related changes in slow wave activity rise time and NREM sleep EEG with and without zolpidem in healthy young and older adults. Sleep Med [Internet]. 2014;15(9):1037–45. Available from: http://dx.doi.org/10.1016/j.sleep.2014.05.007

61. Luca G, Haba Rubio J, Andries D, Tobback N, Vollenweider P, Waeber G, et al. Age and gender variations of sleep in subjects without sleep disorders. Ann Med. 2015;47(6):482–91.

62. Boselli M, Parrino L, Smerieri A, Terzano MG. Effect of Age on EEG Arousals in Normal Sleep. Sleep [Internet]. 1998 Jun 1 [cited 2021 Aug 16];21(4):361–7. Available from: https://academic.oup.com/sleep/article/21/4/361/2731658

63. Landolt HP, Borbély AA. Age-dependent changes in sleep EEG topography. Clin Neurophysiol. 2001 Feb 1;112(2):369–77.

64. Samek W, Binder A, Montavon G, Lapuschkin S, Müller KR. Evaluating the visualization of what a deep neural network has learned. IEEE Trans Neural Networks Learn Syst. 2017 Nov 1;28(11):2660–73.

65. Samek W, Wiegand T, Müller KR. Explainable artificial intelligence: Understanding, visualizing and interpreting deep learning models. arXiv. 2017;

66. Al G, Lan A, L G, Jm H, PCh I, Rg M, et al. PhysioBank, PhysioToolkit, and PhysioNet: Components of a New Research Resource for Complex Physiologic Signals. Circulation [Internet]. 2000;101(23):e215–20. Available from: http://circ.ahajournals.org/content/101/23/e215.full

67. PhysioNet: The Sleep-EDF database [Expanded].

68. Tuk B, Oberyé JJL, Pieters MSM, Schoemaker RC, Kemp B, Van Gerven J, et al. Pharmacodynamics of temazepam in primary insomnia: Assessment of the value of quantitative electrocephalography and saccadic eye movements in predicting improvement of sleep. Clin Pharmacol Ther. 1997;62(4):444–52.

69. Griffin CE, Kaye AM, Rivera Bueno F, Kaye AD. Benzodiazepine pharmacology and central nervous system-mediated effects. Ochsner J. 2013;13(2):214–23.

70. Van Sweden B, Kemp B, Kamphuisen HAC, Van der Velde EA. Alternative electrode placement in (automatic) sleep scoring (F(pz)-C(z)/P(z)-O(z) versus C(4)-A(1)). Sleep. 1990;13(3):279–83.

71. Tsinalis O, Matthews PM, Guo Y, Zafeiriou S. Automatic Sleep Stage Scoring with Single-Channel EEG Using Convolutional Neural Networks. arXiv [Internet]. 2016; Available from: http://arxiv.org/abs/1610.01683

72. Rechtschaffen A, Kales A. A Manual of Standardized Terminology, Techniques and Scoring System for Sleep Stages of Human Subjects. Washington DC: US Government Printing Office; 1968.

73. Youness M. CVxTz/EEG\_classification: v1.0 [Internet]. 2020 [cited 2021 Jan 5]. Available from: https://github.com/CVxTz/EEG_classification

74. Chollet F. Keras [Internet]. GitHub; 2015. Available from: https://github.com/fchollet/keras

75. Abadi M, Barham P, Chen J, Chen Z, Davis A, Dean J, et al. TensorFlow: a system for large -scale machine learning. In: Proceedings of the 12th USENIX Symposium on Operating Systems Design and Implementation. 2016. p. 265–83.

76. Kingma DP, Ba J. Adam: A method for stochastic optimization. arXiv Prepr 14126980. 2014;

77. Sturm I, Lapuschkin S, Samek W, Müller KR. Interpretable deep neural networks for single -trial EEG classification. J Neurosci Methods. 2016 Dec 1;274:141–5.

78. Yan W, Plis S, Calhoun VD, Liu S, Jiang R, Jiang T-Z, et al. Discriminating Schizophrenia From Normal Controls Using Resting State Functional Network Connectivity: A Deep Neural Network and Layer-wise Relevance Propagation Method. In: IEEE INTERNATIONAL WORKSHOP ON MACHINE LEARNING FOR SIGNAL PROCESSING. 2017.

79. Thomas AW, Heekeren HR, Müller K-R, Samek W. Analyzing Neuroimaging Data Through Recurrent Deep Learning Models. 2018 Oct 23; Available from: http://arxiv.org/abs/1810.09945

80. Montavon G, Samek W, Müller KR. Methods for interpreting and understanding deep neural networks. Digit Signal Process A Rev J. 2018;73:1–15.

81. Alber M, Lapuschkin S, Seegerer P, Hägele M, Schütt KT, Montavon G, et al. INNvestigate neural networks! J Mach Learn Res. 2019;20.

